# Temporal Dynamics of Brain Mediation in Predictive Cue-induced Pain Modulation

**DOI:** 10.1101/2024.01.23.576786

**Authors:** Suhwan Gim, Seok-Jun Hong, Elizabeth A. Reynolds Losin, Choong-Wan Woo

## Abstract

Pain is not a mere reflection of noxious input. Rather, it is constructed through the dynamic integration of prior predictions with incoming sensory input. However, the temporal dynamics of the behavioral and neural processes underpinning this integration remain elusive. Here, we identified a series of brain mediators that integrated cue-induced expectations with noxious inputs into ongoing pain predictions using a semicircular scale designed to capture rating trajectories. Temporal mediation analysis revealed that during the early-to-mid stages of integration, the frontoparietal and dorsal attention network regions, such as the lateral prefrontal, premotor, and parietal cortex, mediated the cue effects. Conversely, during the mid-to-late stages of integration, the somatomotor network regions mediated the effects of stimulus intensity, suggesting that the integration occurs along the cortical hierarchy from transmodal to unimodal brain systems. Our findings advance the understanding of how the brain integrates prior and sensory information into pain experience over time.

## Introduction

Pain experience can be understood as a continuous integration process of incoming noxious input with prior contextual information, inferring the current state of both external and internal environments^1–3^. For example, the impact of expectation on pain perception, such as placebo and nocebo effects, can be conceptualized using a Bayesian framework^4, 5^, in which incoming sensory information updates priors, continuously forming new pain priors^6^. Previous studies support this concept, suggesting pain perception as a continuous integration and prediction process across different levels of analysis, from behavioral^7, 8^ and physiological^9^ to neural levels^9, 10^. Moreover, there has recently been an increasing emphasis on examining perception, action, and decision-making as a continuous dynamical process^11, 12^. However, many previous studies have examined brain mechanisms of the integration process by studying each process separately—i.e., pain anticipation, stimulus inputs, and pain ratings—as “individuated elements”^9–11, 13, 14^. To address this gap, our current study investigates the temporal dynamics of brain mediation that underpins the continuous integration process of pain perception.

In this study, we employed temporal mediation analysis to examine the intricate interplay between pain modulatory factors, brain activities, and behavioral outcomes. Previous studies have successfully utilized mediation analysis to identify brain mediators for the pain modulation effects of cue-induced expectancy and the intensity of noxious stimuli^13, 15–17^. For example, the primary and secondary somatosensory cortices (S1 and S2), along with the dorsal posterior insular cortex (dpINS) have been shown to mediate the effects of stimulus intensity on pain perception^16^. The ventrolateral prefrontal cortex (vlPFC) and anterior insular cortex (aINS) have been identified as mediators of cue-induced pain modulation^15^. Furthermore, the frontoparietal and dorsal attention brain networks have been implicated in mediating the effects of social information on pain perception^13^. However, these studies did not focus on the temporal dynamics of the mediation effects, thereby limiting their ability to examine pain as a continuous integration process. To overcome this limitation, we utilized temporal mediation analysis, a method designed to identify time-series of brain mediators throughout the trial.

However, the application of this method faces limitations with conventional experimental paradigms that largely rely on ‘button-based’ or ‘single value-based’ behavioral assessments^9, 10, 13–16^, given that they are typically optimized to probe segmented processes^11^. For example, the traditional pain rating task usually asks participants to report their overall pain intensity or unpleasantness as a single value. This simplification turns the ongoing pain processes into a segmented event, thereby overlooking its inherent dynamic nature. To overcome this limitation, we devised a new rating method, named a semi-circular rating scale, with which we can implement a ‘trajectory-based’ assessment. The ‘trajectory-based’ measurements have proven effective in capturing real-time temporal dynamics of motor or cognitive processes in multiple contexts^12, 18–23^. In our experiment, we asked participants to continuously report their real-time prediction of pain intensity on the semi-circular rating scale using a joystick throughout a trial. We operationalized the rating trajectories as an indicator of a real-time pain integration process^11, 12, 21^. The trajectory-based rating method combined with temporal mediation analysis offers a novel avenue to scrutinize the details of a dynamic process of how the brain integrates sensory inputs with prior contextual information over time.

The primary research questions of the current study are as follows. (1) Can we reveal the continuous integration process underlying pain perception with the trajectory-based measurement? (2) Can we identify the specific timing and brain regions that mediate the temporal dynamics of this integration process? and (3) Are the temporal mediation findings sensible and interpretable in terms of the large-scale functional organization of the brain^24, 25^? We addressed these questions using a functional Magnetic Resonance Imaging (fMRI) experiment using a cue-induced pain modulation task. We employed the semi-circular rating scale to collect trajectory-based ratings of continuous pain prediction. In addition, using this trajectory dynamic information, we conducted temporal mediation analyses to identify brain regions involved in the ongoing integration process. Our hypotheses primarily focus on the potential roles of the lateral prefrontal cortices in encoding contextual information^26–31^. Furthermore, we anticipated that, during this integration process, brain regions across the cortical hierarchy^24, 25^ may exhibit distinct temporal mediation profiles for cue and stimulus effects. For example, transmodal brain regions could play an important role in mediating the effects of contextual cue information, whereas unimodal brain regions may primarily mediate the stimulus effects on pain. Overall, this study seeks to elucidate when and which brain regions and networks contribute to pain construction at different temporal stages.

## Results

### Behavioral results

Inside the scanner, participants performed a cue-induced pain modulation task involving continuous pain prediction ratings. Participants were instructed to continuously report their ongoing pain predictions in response to the question, “How much pain do you predict?” (**Fig. 1a**) Using the semi-circular rating scale, we obtained participants’ ongoing pain predictions and instructed them to treat the angle as the level of pain regardless of the distance from the initial point, i.e., the center of the rating scale. Prior to heat stimulation, 25 dots appeared on the edge of the semi-circular rating scale. Participants were informed that the dots represented other participants’ pain ratings from previous experiments about the upcoming stimulus^13, 32^. These dots served as pain-predictive cues based on social information. In practice, we displayed 25 dots randomly sampled from a normal distribution for high pain cue (mean: 138.6° sd: 9°), low pain cue (mean: 39.6°; sd: 9°), and no cue condition^8^ (**Fig. 1b**). After the cue presentation and jittered inter-stimulus interval, participants experienced five different levels of 12.5 seconds heat stimulation while reporting their continuous pain prediction ratings. The five levels of stimulus temperature were fitted for each participant through a calibration procedure prior to the fMRI scans^15^ to provide similar levels of pain intensities across participants (see **Supplementary Fig. 1** for the calibration results and **Methods** for the detailed experimental procedure). The continuous ratings were collected for 14.5 seconds, including 1 second no heat period before and after the heat stimulation. After the heat stimulation, we additionally asked participants to rate their overall pain intensity rating for the stimulation.

**Fig. 1.**
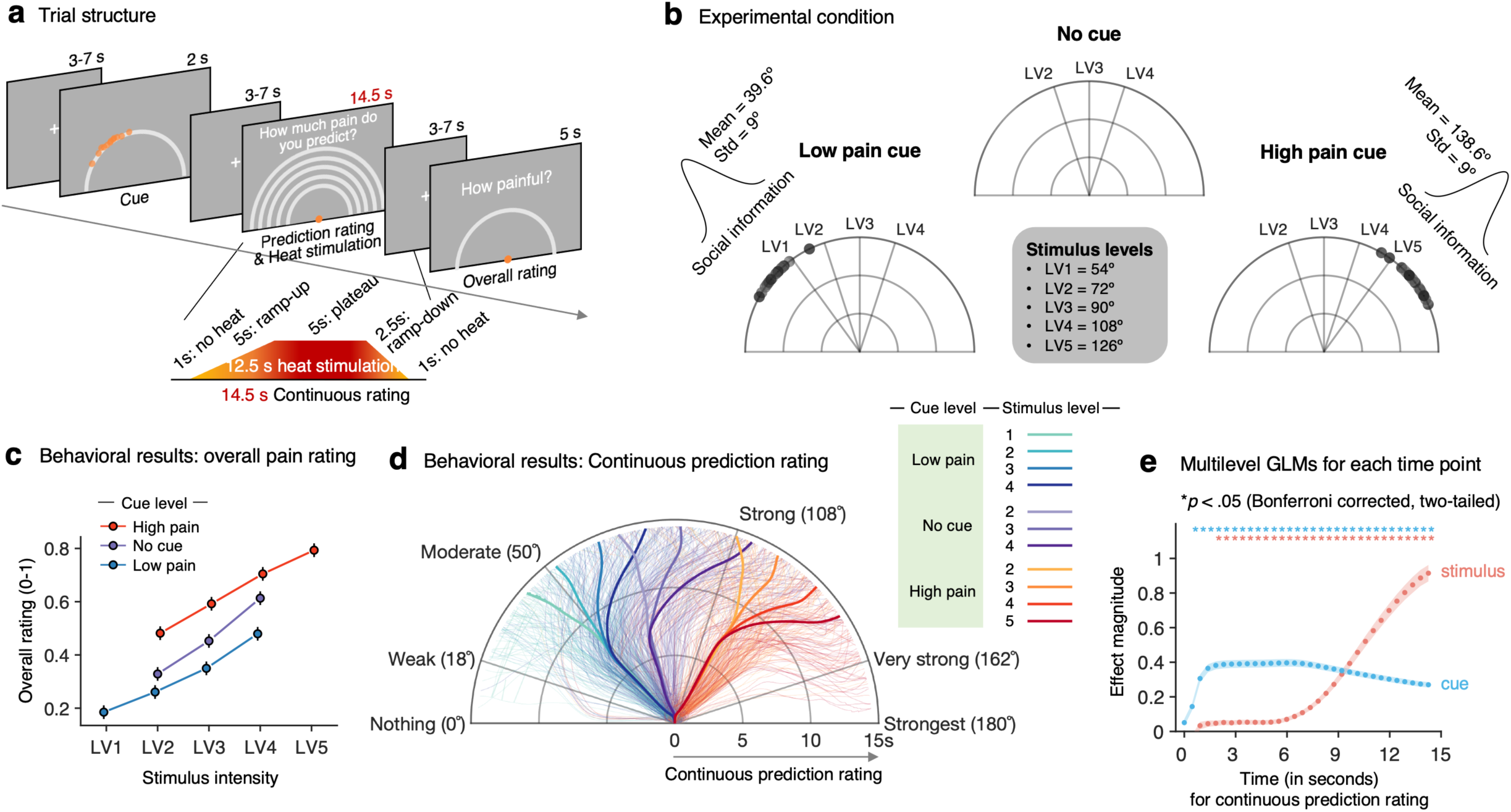
Experimental paradigm and behavioral results. **a,** Trial structure: Each trial started with a cue phase, except for the ‘No cue’ trials, which began with thermal stimulation. Participants were instructed to continuously report their ratings in response to the following question, “How much pain do you predict?” At the end of each trial, participants provided their overall pain rating. **b,** Experimental conditions: There were three cue conditions: ‘low pain cue,’ ‘no cue,’ and ‘high pain cue.’ Cues consisted of twenty-five dots randomly sampled from a normal distribution with varying means and a standard deviation. Participants were informed that the dots represented other participants’ pain ratings for the upcoming stimulus. We delivered heat stimuli at five temperature levels calibrated for each participant to induce similar levels of pain intensity. **c,** Overall pain ratings: The plot shows the average overall pain ratings as a function of stimulus intensity. Different colors represent different cue conditions. Error bars represent the within-subject standard errors of the mean (s.e.m.). **d,** Continuous prediction ratings: Displayed are the trajectories of pain prediction ratings. Thick lines represent the group-average trajectories, while thin lines show the individual-level averages of trajectories. Different colors indicate different experimental conditions. **e,** Multilevel GLM results: We conducted a series of multilevel GLM analyses, in which we included pain prediction ratings at each time point as a dependent variable and the cue and stimulus intensity conditions and their interaction as independent variables. Given that the continuous ratings were segmented into 32 bins, a total of 32 multilevel GLM analyses were performed. To correct for multiple comparisons, we applied the Bonferroni correction method. Within-subject s.e.m. is denoted by shaded error bands in the plot. Asterisks above the plot indicate time points with significant effects at *p* < 0.05, Bonferroni corrected, two-tailed. The plot uses blue to denote cue effects and red for stimulus effects.

To assess the effects of the pain-predictive cues and the stimulus intensity on overall pain ratings, we performed a multilevel general linear model (GLM) analysis. The model predicted the overall pain intensity ratings using cue conditions and stimulus intensity as independent variables. We found significant main effects of both the cue level (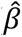 = 0.109, 95% confidence interval (CI) = [0.094, 0.125], *z* = 3.836, *p* < 1.2479 × 10^-4^, bootstrap test with 10,000 iterations) and the stimulus intensity level (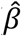 = 0.106, 95% CI = [0.096, 0.117], *z* = 3.724, *p* < 1.9549 × 10^-4^, bootstrap test) (**Fig. 1c**), suggesting that the manipulation of contextual information substantially influenced pain perception^8, 13, 32^.

Next, to investigate the temporal dynamics of the cue and stimulus effects on the continuous ratings, we first resampled the ratings at 50Hz using linear interpolation. We then averaged the ratings over 32 data points, ensuring each time bin matched the fMRI sampling rate, TR = 0.46 seconds. The rating angle, which ranged from 0° to 180°, was transformed to ratings on a scale from 0 to 1. The average trajectory patterns across different conditions indicated that the initial rating direction aligned with the cue location. This direction persisted until roughly 7 seconds, coinciding with the plateau phase of the heat stimulation (**Fig. 1d** and **Supplementary Fig. 2**). After the plateau, the ratings began to gravitate toward the stimulus intensity level, suggesting that while the cue effects were predominant in the early phase, the stimulus intensity effects became important in the later phase. These observations were corroborated by a series of multilevel GLMs to model the trial-by-trial variations in ratings at each time point based on the levels of cues and stimulus intensity^21, 33^. As shown in **Fig. 1e**, both the cues and the stimulus intensity significantly influenced the continuous pain prediction ratings throughout the trial, with the exception of stimulus effects being non-significant during the initial 1.4 seconds, which included the 1-second no-heat period (*p*s < 0.05, Bonferroni corrected, two-tailed). Examining the magnitude of the effects, the cue had a larger effect than the stimulus intensity up to 9 seconds, with its impact gradually decreasing over time after that. Conversely, the influence of the stimulus intensity increased over time, surpassing the cue effects after around 9 seconds. In summary, our behavioral analysis showed that the cue and stimulus effects exhibited distinct temporal profiles, revealing a gradual shift in their relative importance from contextual to sensory information integration.

### Brain mediators of the cue and stimulus effects on the overall pain ratings

Initially, as done in previous studies^13, 15–17^, we first sought to identify the brain areas mediating the effects of the cue and stimulus intensity on the overall pain ratings before exploring the detailed temporal dynamics of the integration process. To search for brain mediators, we performed a whole-brain search using multilevel mediation analysis ^15, 17^ (**Fig. 2a** and **2b**, left). In this analysis, the trial-by-trial experimental condition, such as cue level or stimulus intensity, was included as a predictor (*x*), overall pain rating as an outcome (*y*), and single-trial brain activation maps during the heat stimulation period as mediators (*m*). When cue condition served as a predictor, stimulus intensity was included as a covariate, and in the model where the stimulus intensity served as a predictor, the cue levels were included as a covariate. The findings showed multiple significant brain mediators for the cue effects at *q* < 0.05, false discovery rate (FDR) corrected, including the anterior insula (aINS), supplementary motor area (SMA), temporal pole (TP), vlPFC, and visual and somatomotor cortex (SMC) areas (**Fig. 2a**). Notably, the vlPFC, aINS, and visual cortex regions have been previously associated with cue-induced pain modulation ^13, 15^. For the effects of stimulus intensity, significant brain mediators at FDR *q* < 0.05 featured some regions overlapping with those identified for cue mediation, including the SMA, SMC, aINS, and visual cortex (**Supplementary Fig. 3**). However, unique brain mediators distinct from those in cue mediation were also identified, including the dpINS and thalamus. In addition, there was a more pronounced involvement of the SMC areas, including the S2 (**Fig. 2b**). These regions have also been previously linked to pain processing in the brain^16, 34–37^. Overall, the results of the conventional mediation analysis for the overall pain ratings were largely consistent with findings from previous studies.

**Fig. 2.**
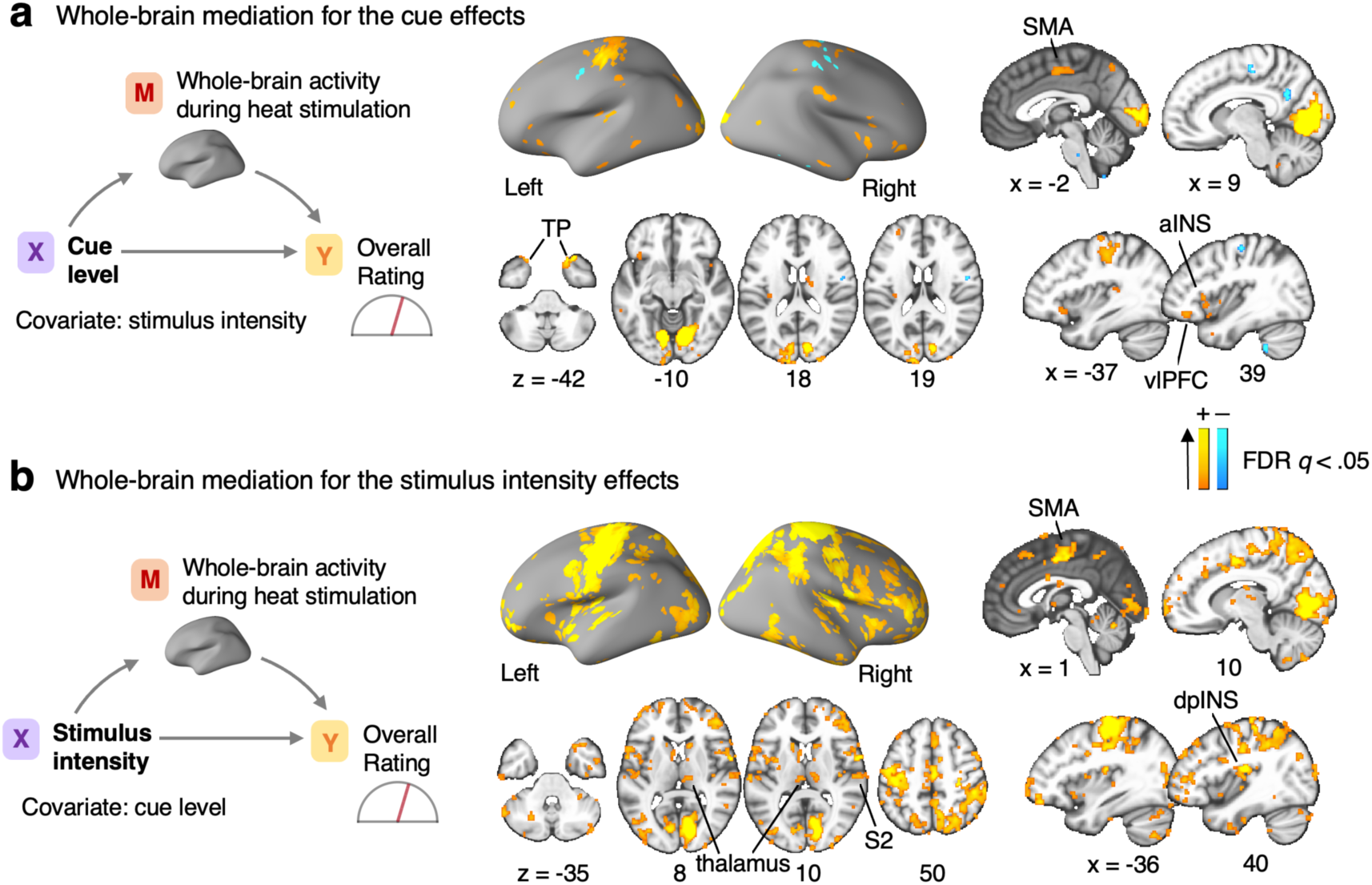
Brain mediators of cue and stimulus intensity effects on overall pain ratings. **a,** The mediation model assessed how cue levels influenced overall pain ratings, with single-trial brain activation during heat stimulation as the mediator and stimulus intensity as a covariate. The brain map displays significant brain mediators for the cue effects. **b,** The mediation model assessed how stimulus intensity influenced overall pain ratings, with brain activation as the mediator and cue level as a covariate. The brain maps were thresholded using a false discovery rate (FDR) of *q* < 0.05 and a minimum cluster size of k > 5 voxels. SMA, supplemental motor area; aINS, anterior insular cortex; dpINS, dorsal posterior insular cortex; vlPFC, ventrolateral prefrontal cortex; TP, temporal pole.

### Temporal dynamics of brain mediation for cue and stimulus effects

Finally, to investigate the dynamical integration process of cue and stimulus information in the brain, we analyzed the temporal dynamics of brain mediation for the cue and stimulus effects on the continuous pain prediction ratings. To this end, we performed the whole-brain multilevel mediation analysis with combinations of 45 TR (= 20.7 seconds) heat-evoked brain activity from the single-trial finite impulse response (FIR) model and 32 segmented continuous pain prediction ratings (= 14.5 seconds) (**Fig. 3a**; see *Methods* for details). We included a longer duration for the brain activation maps to accommodate potential hemodynamic response delays. To help interpretation of the results, we segmented the 20.7-second brain activation period into three time domains—early (0-6 s), middle (6-13.8 s), and late (13.8-20.7 s). We also divided the 14.5-second rating period into the following time domains—cue dominant (1-6 s), transition (6-10 s), and stimulus dominant (10-14.5 s) based on the multilevel GLM results shown in **Fig. 1e**. In addition, to define time domains for identifying significant brain mediators, we employed a data-driven approach via independent component analysis (ICA)^38^. This method allowed us to identify significant and interpretable spatiotemporal domains for brain mediation. As illustrated in **Supplementary Fig. 4**, we derived 5 independent components (ICs) and first determined interpretable time domains for mediation based on the top 2.5 percentile of the temporal weights. This thresholding level was somewhat arbitrary, but it yielded reasonable and interpretable time windows. These time domains represented the temporal connection between the specific time periods of the heat-evoked brain activity and the continuous pain prediction ratings (see **Supplementary Fig. 4a**). We then identified the brain mediators for each time domain based on pre-established criteria for selecting brain mediators. Specifically, we selected voxels that covered at least 5 percent of the defined time domain and survived at FDR *q* < 0.05 with a cluster size greater than 5 voxels.

**Fig. 3.**
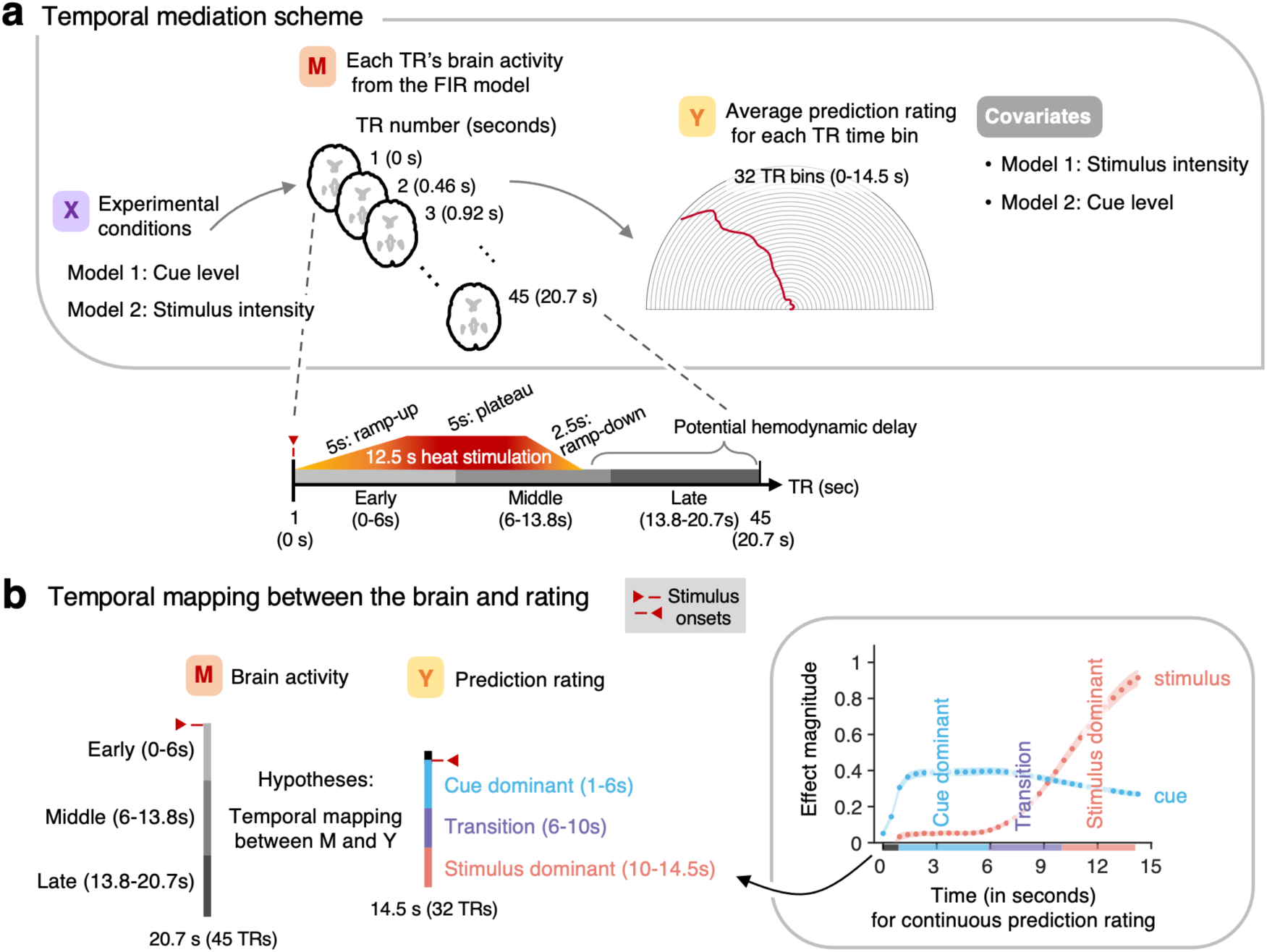
Temporal mediation analysis. **a,** A schematic representation of whole-brain multilevel mediation analysis with 45 TR heat-evoked brain activation patterns from a single-trial finite impulse response (FIR) model and 32 time-segmented pain prediction ratings. The temporal composition of the 45 TRs is shown below. The temporal mediation analysis quantified the influence of experimental conditions (*x*) on brain activity (*m*), and how these brain activity patterns relate to continuous pain prediction ratings (*y*) accounting for the experimental conditions at each time bin. In the stimulus intensity model, cue level served as a covariate, and conversely, stimulus intensity was a covariate in the cue mediation model. **b.** To help interpretation of the temporal mediation results, we divided the brain activity into three time domains—early (0-6 s), middle (6-13.8 s), and late (13.8-20.7 s). We also divided the 14.5-second rating period into the following time domains—cue dominant (1-6 s), transition (6-10 s), and stimulus dominant (10-14.5 s) based on the multilevel GLM results.

Through the temporal mediation analysis of the cue effects, we identified four main time domains (as illustrated in **Fig. 4a** and **Supplementary Fig. 4b**). These time domains included the mapping between the early and middle phases of heat-evoked brain activity and the cue dominant and transition phases of the continuous pain prediction ratings, suggesting that the early and middle phases of the brain activation are important for mediating the cue effects. The supra-threshold brain regions within these four temporal domains included the dorso- and ventro-lateral PFC (dlPFC and vlPFC), intraparietal sulcus (IPS), TP, and visual cortex areas that have been associated with pain predictive cue-induced pain modulation in previous studies ^13–15^ (**Figs. 4b-c**). Regarding the large-scale functional networks, the brain regions associated with early mediation of cue effects engaged the dorsal and ventral attention and frontoparietal networks, and their mediation effects waned over time. Conversely, the mediation of cue effects of the somatomotor (particularly in the left hemisphere) and visual networks increased over time. This suggests a transition in the brain systems’ engagement from higher-order cognitive processes to visuomotor functions in relation to cue effects. This idea is further supported by the analysis using the principal gradient map ^24^ derived from the resting-state fMRI data of the participants in this study (**Fig. 4d**). When we calculated the proportions of different levels of principal gradients for the four thresholded brain mediation maps ^39, 40^, the temporal dynamic patterns clearly showed that the transition from transmodal to unimodal brain areas in mediation cue effects.

**Fig. 4.**
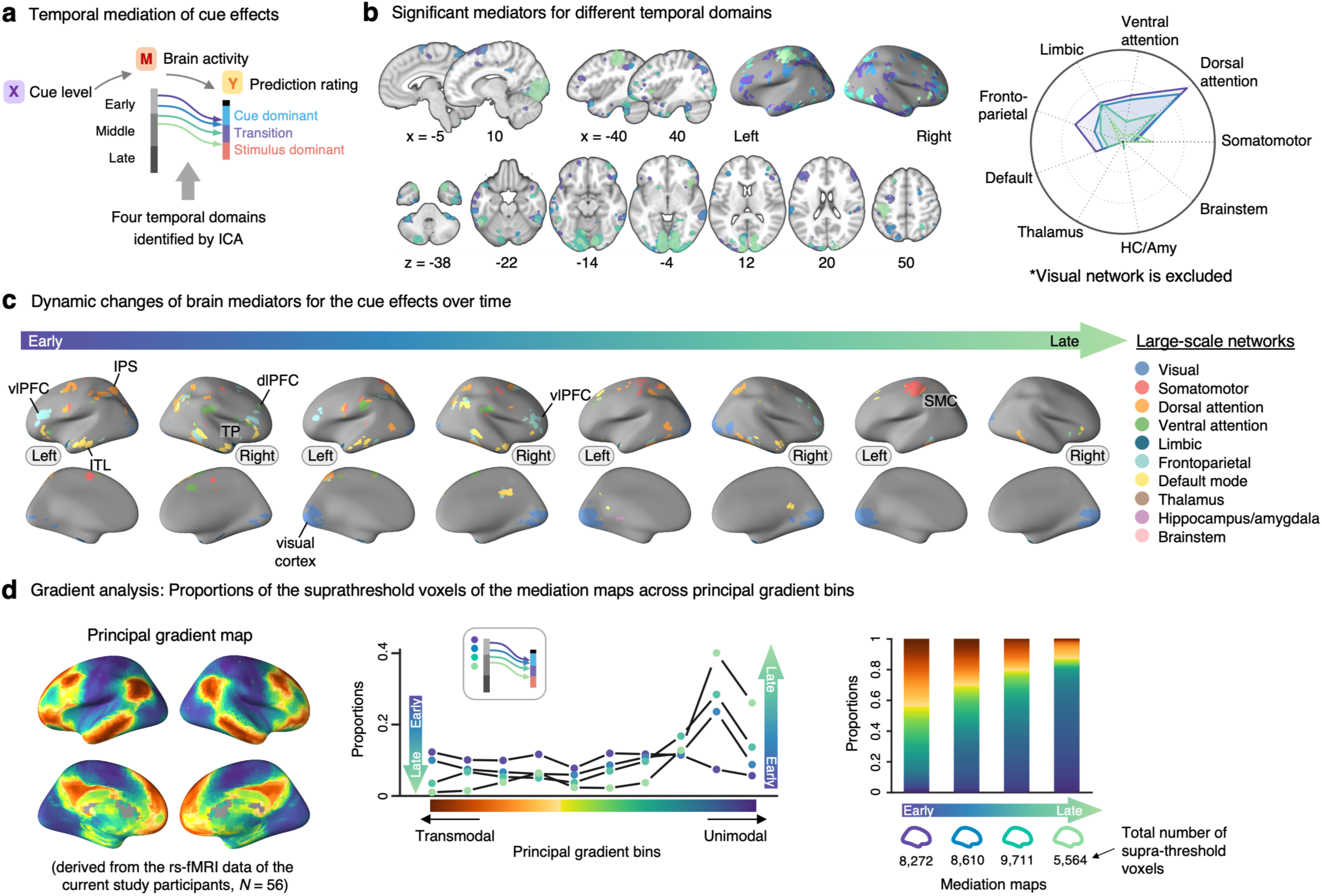
Temporal mediation analysis results for the cue effects. **a,** A schematic diagram showing four temporal domains identified with the ICA (see **Supplementary Fig. 4**) in the cue mediation model. Lines with different colors represent the mapping between the early and middle phases of heat-evoked brain activity and the cue dominant and transition phases of the continuous pain prediction ratings. **b,** The brain map displays the supra-threshold brain mediators for the cue effects. The four distinct colors in this map correspond to the four temporal domains. The radial plot shows the relative proportions of the number of overlapping voxels between the thresholded mediation maps and each network (or region), given the total number of voxels within each network (or region). We did not include the visual network in the radial plot. It is because the visual cortex activation would mainly reflect the task-related features, but it dominated the plot, making it difficult to interpret the activation patterns over time. **c,** The brain mediation maps corresponding to the four temporal domains are color-coded based on large-scale functional and subcortical brain networks. **d,** (Left) The volumetric functional gradient map was derived from resting-state fMRI data of the current study participants (*N* = 56). (Middle, Right) We divided the principal gradient into two sets of bins: one set comprising 10 bins using 10-percentile intervals (Middle) and another set of 100 bins using 1-percentile intervals (Right), creating binary gradient bin masks for each set. We then calculated the proportions of overlap between the thresholded brain maps and these gradient bin masks. In the middle panel, the plot shows the overlaps between four mediation maps corresponding to four temporal domains and 10-bin principal gradient masks. Mediation maps are represented with different dot colors. The gradient transitioning from red to blue represents the principal gradient spectrum, ranging from transmodal to unimodal brain areas. In the right panel, we visualized the overlaps between the same four mediation maps and 100-bin principal gradient masks using a stacked bar chart. Abbreviations: HC/Amy, hippocampus and amygdala; SMC, somatomotor cortex; dlPFC, dorsolateral prefrontal cortex; vlPFC, ventrolateral prefrontal cortex; TP, temporal pole; ITL, inferior temporal lobe; IPS, intraparietal sulcus.

We next identified four main time domains and brain mediators of the stimulus effects (**Fig. 5a**). These time domains showed the mapping between the middle-to-late phases of brain activity and the stimulus-dominant phase of ratings. The supra-threshold brain regions within these time domains also included the dl/vlPFC, IPS, TP, and visual cortex areas as in the cue mediation maps, but the stimulus mediation maps additionally showed strong brain mediations in the somatomotor network regions, such as the SMA, dpINS/S2, and primary sensory and motor cortex (SMC) areas (**Figs. 5b-c**). Importantly, the total number of significant voxels increased over time, suggesting that the stimulus mediation effects intensified over time, particularly at the later stages of integration. In terms of the large-scale functional networks, brain regions within the somatomotor and visual networks showed heightened mediation effects in the later phase. In addition, the dorsal attention and frontoparietal network regions also demonstrated increased involvement during the later phase of integration. Additional analyses with the principal gradient map showed a similar pattern to the cue mediation maps—i.e., the transition from transmodal to unimodal brain areas over time was observed in the stimulus mediation maps (**Fig. 5d**).

**Fig. 5.**
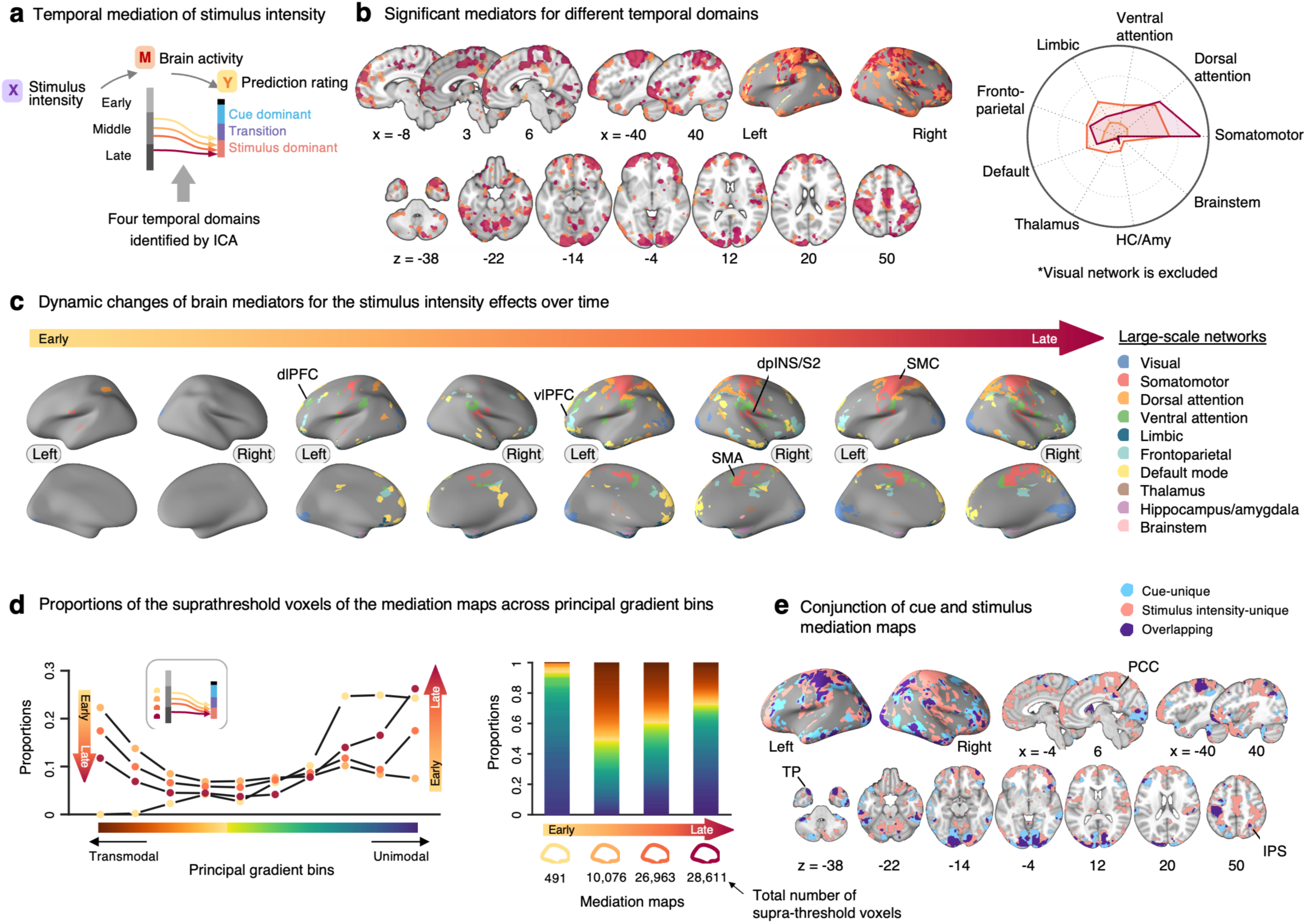
Temporal mediation analysis results for the stimulus effects. **a,** A schematic diagram showing four temporal domains identified with the ICA (see **Supplementary Fig. 4**) in the stimulus intensity mediation model. Lines with different colors represent the mapping between the middle and late phases of heat-evoked brain activity and the stimulus-dominant phase of the continuous pain prediction ratings. **b,** The brain map displays the supra-threshold brain mediators for the effects of stimulus intensity. The four distinct colors in this map correspond to the four temporal domains. The radial plot shows the relative proportions of the number of overlapping voxels between the thresholded mediation maps and each network (or region), given the total number of voxels within each network (or region). We excluded the visual network on radial plot because it reflected task characteristics, making it difficult to understand the patterns over time. **c,** The brain mediation maps corresponding to the four temporal domains are color-coded based on large-scale functional and subcortical brain networks. **d,** (Left, Middle) Similar to Fig. 4d, we displayed the proportions of overlap between thresholded brain maps and the gradient bin masks. In the left panel, the plot shows the overlaps between four mediation maps corresponding to four temporal domains and 10-bin principal gradient masks. Mediation maps are represented with different dot colors. The gradient transitioning from red to blue represents the principal gradient spectrum, ranging from transmodal to unimodal brain areas. In the middle panel, we visualized the overlaps between the same four mediation maps and 100-bin principal gradient masks using a stacked bar chart. **e,** The conjunction map shows the spatial overlap and unique areas involved in the cue and stimulus mediation. Regions colored in cyan and pink denote brain areas uniquely associated with the mediation of cue and stimulus effects, respectively. The areas where these two mediation maps overlap are colored in purple. Abbreviations: HC/Amy, hippocampus and amygdala; SMC, somatomotor cortex; dlPFC, dorsolateral prefrontal cortex; vlPFC, ventrolateral prefrontal cortex; dpINS, dorsal posterior insular cortex; S2, secondary somatosensory cortex.

When we compared the cue and stimulus mediation maps, there were multiple overlapping brain regions (**Fig. 5e**). For example, motor and visual cortex regions appeared to mediate both cue and stimulus effects, which may be related to the task-related factors, such as operating a rating apparatus and looking at the rating trajectory on the screen. In addition to these regions, there were other overlapping brain mediators, such as subregions within the vlPFC, IPS, TP, inferior temporal lobe (ITL), and posterior cingulate cortex (PCC), which may play a role in maintaining the cue information throughout the whole integration period, facilitating the integration process of the cue and stimulus information. There were also brain regions that exclusively contributed to the mediation of either the cue or stimulus effects. For example, brain regions within the dorsal and ventral attention networks, including dlPFC and aINS, were among the unique brain mediators for the cue effects. For the stimulus effects, multiple brain regions within the somatomotor network, such as the dpINS, SMA, and right SMC regions, appeared as unique brain mediators. Notably, the anterior and posterior parts of the insular cortex differentially mediated the cue and stimulus effects, respectively, which was consistent with the mediation analysis results on the overall pain ratings as shown in **Supplementary Fig. 3** and previous studies ^9, 41, 42^. To summarize the temporal dynamics of the overlapping brain regions’ mediation effects, selected regions’ temporal mediation profiles were visualized in **Fig. 6**. The results show that many overlapping brain mediators showed significant mediation effects during the early phase of the cue effects and the late phase of the stimulus effects, suggesting their role in representing prior and posterior information of pain.

**Fig. 6.**
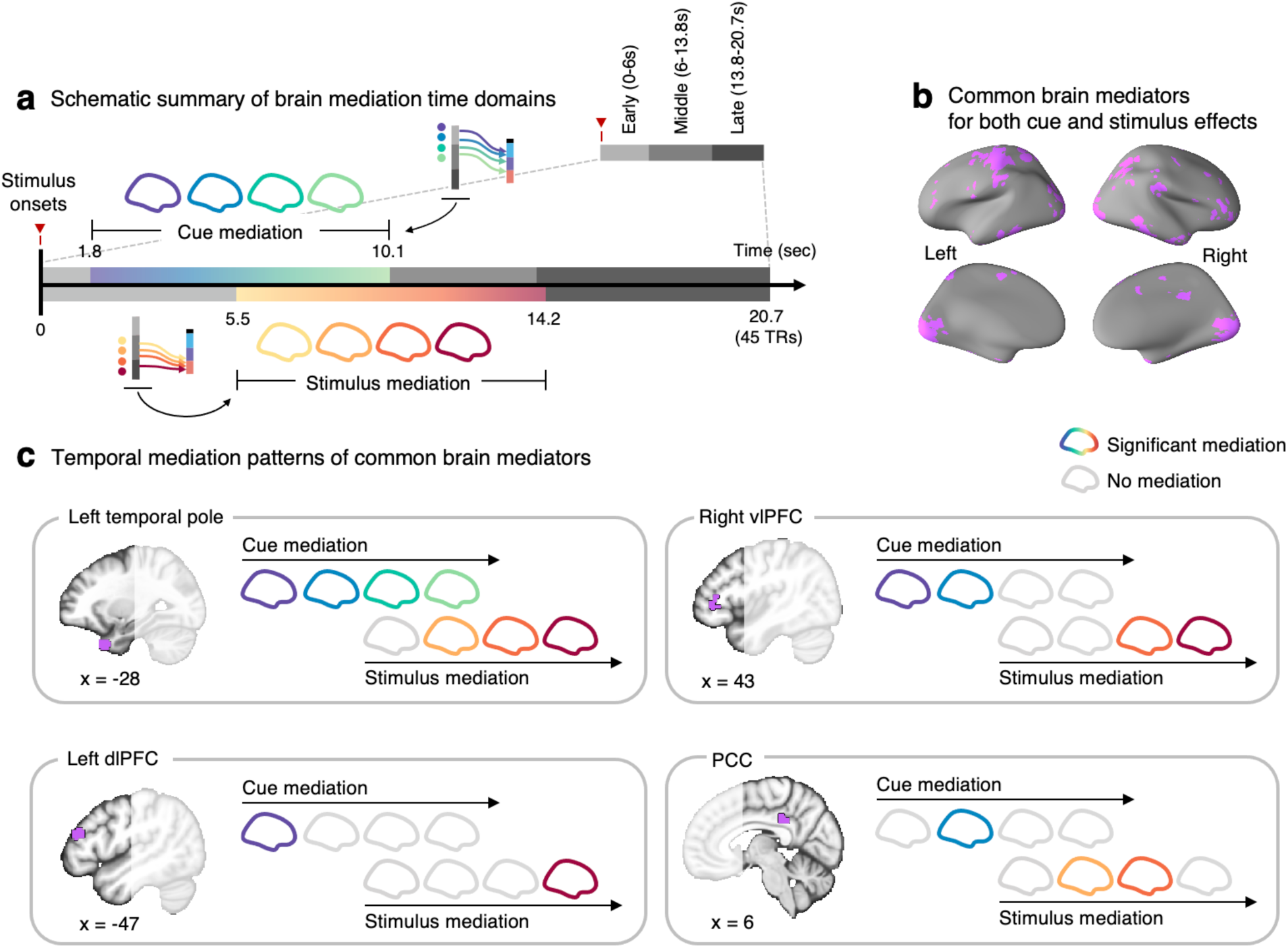
Temporal mediation patterns of shared brain mediators. **a,** The schematic serves as a visual summary, depicting the brain mediation time domains identified through independent component analysis (ICA). The duration of mediation effects—cue mediation from 1.8 to 10.1 seconds post-stimulus onset, and stimulus mediation from 5.5 to 14.2 seconds—is marked on the timeline based on the ICA results. **b,** The brain map shows common brain mediators (cluster extent *k* > 2) for both cue and stimulus effects. **c,** The specific time domains where selected common brain mediators exhibit significant mediation effects. Colored brains correspond to periods of significant mediation, whereas gray indicates non-significant mediation. Abbreviations: dlPFC, dorsolateral prefrontal cortex; vlPFC, ventrolateral prefrontal cortex; PCC, posterior cingulate cortex.

## Discussion

The current study investigated the temporal dynamics of brain mediation in predictive cue-induced pain modulation. Our findings provide comprehensive temporal profiles of behavioral and neural processes underlying the complex interplay between cognitive and sensory components of pain ^25, 43^. The pain experience can be conceptualized as a continuous Bayesian updating process, in which predictions about the pain state are continuously updated by integrating contextual information with incoming sensory inputs. Particularly in this study, we examined the temporal dynamics of brain mediation underlying this continuous integration process for pain by utilizing novel behavioral and analysis methodologies, including a semicircular pain rating scale and a temporal mediation analysis method.

In this study, we introduced a semi-circular rating scale, presenting a new method of conducting trajectory-based analyses on rating behaviors to quantitatively describe and visualize the temporal dynamics of pain perception. A common challenge in utilizing online ratings in fMRI studies with the conventional linear rating scale lies in the potential confounding effects arising from the correlation between rating-related motor responses and pain intensity^44^. For example, to indicate higher pain intensity, participants are required to move the cursor further to the right or upwards^45, 46^. In the semi-circular rating scale, the use of angle rather than distance to indicate the level of pain allows us to minimize the impact of movement-related confounds. In addition, the trajectory information derived from pain ratings can reveal cognitive states through the quantification of ongoing cognitive processes that would otherwise remain hidden^12, 18^. For example, previous studies have demonstrated that the rating trajectory can provide information about hidden cognitive dynamics, such as number comparison^19^, food choice influenced by health-related cues^21^, and face perception influenced by stereotypes^20^. Unlike button press-based assessments that primarily provide reaction time or performance data, trajectory-based measurements can offer more detailed and richer information about the spatiotemporal patterns of ongoing cognitive processes, which may vary across contexts and among individuals.

Furthermore, we implemented temporal mediation analyses to more effectively model the ongoing effects of cues and stimuli on the brain and the rating trajectory. This analysis method allows us to go beyond conventional mediation analysis approaches ^13, 15, 17^, which typically focus on a snapshot of the brain-behavior relationship. The temporal mediation approach can elucidate the temporal dynamics of the brain-behavior relationship. In addition, we adopted a data-driven approach to identify distinct temporal domains important for the cue and stimulus effects on the brain, specifically utilizing Independent Component Analysis (ICA)^38^. The integration of the temporal mediation methods with ICA offers a new quantitative way to explore the temporal dynamics of the brain-behavior relationship.

The analysis of the rating trajectories highlighted the gradual changes in the effect magnitudes of both the cue and the stimulus—from a cue-dominant phase through a transition phase to a stimulus-dominant phase. This finding, although could potentially offer an important mechanistic perspective to pain perception, has not been clearly demonstrated in previous research. Prior studies have mainly focused on the overall effects of cue and stimulus intensity on pain ratings, often overlooking how these effects unfold over time ^8, 13, 15, 47, 48^. The current study, however, views pain perception as a continuous dynamic process^11^, in which incoming sensory information continuously updates priors, thereby shaping the pain experience^6, 49, 50^. An additional analysis correlating continuous pain prediction ratings with overall post-stimulus pain ratings revealed a gradual increase in similarity over time (**Supplementary Fig. 5**), demonstrating the temporal progression in updating priors for the dynamic construction of the pain experience.

The results from our temporal mediation analyses indicate that brain regions within the frontoparietal and dorsal attention networks, such as the dl/vlPFC and IPS, were among key brain mediators of the cue effects during the early time domains. This aligns with previous studies indicating the involvement of these networks in sensory integration and the attentional modulation of pain ^51–53^. Moreover, they have been identified as mediators in the context of social influences on pain^13^. Further examination of the signs of paths *a* and *b* for the cue mediation effects (**Supplementary Fig. 6a**) revealed that most regions within the frontoparietal and dorsal attention networks showed positive associations with cue levels and pain prediction ratings, highlighting their facilitating role in cue-induced pain modulation^51, 52^. For the stimulus intensity effects, brain regions within the somatomotor network, such as the SMA, SMC, and dpINS/S2 areas, appeared to be the key brain mediators. The mediation effects of these regions became stronger over time during the middle-to-late phases. These findings are consistent with earlier research suggesting the significance of these brain regions in representing subjective pain intensity^35, 37, 54–57^ and mediating the effects of stimulus intensity^16, 17^.

Importantly, there were also overlapping brain mediators between the cue and stimulus effects, including the dl/vlPFC, IPS, TP, ITL, and PCC, as shown in **Fig. 5e**. Many of these regions demonstrated significant mediation effects spanning the early-to-late periods of the trial (**Fig. 6c**). Prior studies have highlighted the crucial roles of the lateral prefrontal cortices, specifically the dlPFC and vlPFC, in encoding contextual information and filtering out irreverent details^26–31, 49^. Notably, the dl/vlPFC have been implicated in mediating the effects of cue and stimulus intensity on pain ^15, 16^. Additionally, the dlPFC, in conjunction with the TP, has been suggested to shape the affective tone related to short-term memories of pain experience^58^, and distinct temporal profiles of the TP and vlPFC have been linked to working memory processes^59^. Together, these findings suggest that the overlapping brain mediators, particularly dl/vlPFC and TP may play crucial roles in maintaining contextual and sensory information as a central integration hub, assembling evidence over an extended timeframe to construct the subjective experience of pain.

Furthermore, the temporal mediation results of our study can be interpreted within the context of a cortical hierarchy as proposed by Mesulam^25^ and further elaborated by Margulies et al. ^24^. Mesulam’s framework (1998) proposes a hierarchical organization of the brain that supports the progression of information processing from sensory inputs to higher cognitive functions ^25^. This hierarchical organization differentiates between unimodal brain regions, which receive sensory inputs from single sensory modalities, and transmodal brain regions involved in multimodal representation and top-down modulations. Our findings suggest that the integration of cue and stimulus information in pain processing involves a dynamic transition from transmodal to unimodal brain regions. The process begins with the encoding of pain predictions in transmodal brain regions, followed by the processing and integration of stimulus information in unimodal brain regions. To the best of our knowledge, our study offers the first demonstration of the integration of pain prediction into pain perception being represented along such a cortical hierarchy.

The temporal mediation analyses in our study, while informative, are not without limitations. First, determining temporal domains and thresholding the spatial maps was somewhat arbitrary. Second, we had to reduce voxel size in heat-evoked brain images from 2mm^3^ to 4mm^3^ to alleviate computational load. Third, the use of five components for estimating temporal domains via ICA could have resulted in missing other components that might be important and have distinct temporal patterns. To address these limitations, we additionally tested seventeen pre-defined ROIs (**Supplementary Fig. 7a**) known to be associated with pain processing ^60^, self-regulatory strategies ^17, 61^, and pain expectation manipulated by social information ^13^. The results suggested that many brain regions showed similar temporal mediation patterns as those identified in whole-brain temporal mediation analysis, including the vlPFC, vmPFC, OFC, and IPS as mediators of cue effects on pain, and the cerebellum, pgACC, SMG, thalamus, dpINS, vlPFC, and IPS as mediators of stimulus intensity effects on pain (**Supplementary Figs. 7b** and **7c**). However, it also revealed unique temporal mediation patterns, as described in **Supplementary Fig. 7**. To expand the current findings and explore additional spatiotemporal patterns of temporal domains, future studies could incorporate a greater number of components and higher spatial resolutions.

The continuous pain prediction task also has some limitations. Most importantly, it was inherently susceptible to head and arm movements due to the motion required for reporting their pain prediction and pain ratings using the MR-compatible joystick. To mitigate the influence of movements on heat-related BOLD signals, we implemented two strategies. First, we increased the number of participants, 59 participants, in comparison to previous studies: 38^13^, 30 (^10^, study1), and 34 (^10^, study2) participants. It allows us to reduce the possibility that the current results might be influenced by a small number of outliers with extreme head and/or arm movements. Second, we excluded the single-trial heat-evoked estimates with 2.5 variance inflation factors (VIF) as high VIFs indicate potential multicollinearity between head movement and effects of interest. Moreover, excluding high VIFs (> 2.5) can minimize the effects of non-interest such as signals of cerebrospinal fluid and white matter, leading to more reliable results. The average number of excluded trials was 3.51 (about 3%; *SD* = 2.09) of the total 108 trials per participant. As such, we believe our results are unlikely to be significantly affected by movement artifacts. Future studies would benefit from methods that can alleviate the impact of non-BOLD artifacts, such as multi-echo fMRI ^62, 63^.

Overall, the current study provides new insights into the temporal dynamics of pain perception and enhances our comprehension of underlying neural mechanisms. Utilizing novel tools and analysis approaches, our results support the notion that pain perception is a dynamic process of continuously integrating contextual information with sensory inputs. This underscores the importance of investigating the temporal dimension of pain perception. We hope that our findings will contribute to a more profound understanding of the brain mechanisms responsible for shaping and modulating the experience of pain.

## Methods

### Participants

We recruited a total of 84 healthy participants who were right-handed, had no neurological disorders, and agreed to participate in a pain experiment. Among them, 59 participants (Korean native speakers; *n*_female_ = 26; mean_age_ = 22.11; range_age_ = 18-28; SD_age_ = 2.44) completed the experiment. Specifically, 84 participants came to the first session that included the pain calibration task ^8, 15^ and a battery of individual difference questionnaires (e.g., demographic, psychological status, and health questionnaires). Thirteen did not continue the experiment because their pain calibration results exceeded the exclusion criteria and eight participants notified withdrawal from the next session due to their own personal reasons. Thus, remaining 63 participants underwent the fMRI task. Among them, three participants requested to quit the experiment during the scan, and one participant showed pain ratings substantially different from the pain calibration results. Participants were recruited through the university websites and flyers posted on the university buildings. All participants were asked to write an informed consent before the experiment in both the first and second sessions and received a monetary reward for their participation. The experiment was approved by the institutional review board at Sungkyunkwan University. The present study collected data from spring to autumn 2018.

### Thermal stimulation

Thermal stimulation that ranged from 40 to 49.2°C (baseline: 32°C) was delivered to a left forearm using a 16 × 16 mm ATS thermode (Medoc, Israel). The thermal stimulation had a duration of 12.5 seconds, comprising a 5-second ramp-up, a 5-second plateau, and a 2.5-second ramp-down (**Fig. 1a**). During the pain calibration task, we changed the stimulation sites on the forearm for each trial, and similarly, for each experimental run during the fMRI task. Four skin sites were used for the former and three for the latter. A single highest temperature of heat stimulation was delivered before the pain calibration task (e.g., 49.2 °C) and each run during the fMRI experiment (e.g., temperature corresponding to the stimulus intensity level 5 from the result of the pain calibration task) to avoid the initial habituation of the skin site to contact heat ^64^ and to confirm the working of heat stimulation delivery.

### Semicircular rating scale

To collect continuous pain prediction ratings, we devised a new rating scale, named the semicircular rating scale. For continuous ratings, we made the scale surrounded by multiple semi-circles to imply that it is a continuous field, while a single semicircle was used for the overall rating (**Supplementary Fig. 8**). An important feature of the semicircular rating scale is that the starting point is equidistant from all possible ratings, addressing an issue of intensity rating being confounded with the cursor movement distance. An orange-colored dot was located at the center of the scale, and the dot can be operated by input devices such as a joystick, which we used in this study. Participants were asked to report their ratings using the angle from the left

segment of the semicircle base by manipulating the orange-colored dot (see **Supplementary Fig. 8**). The rating trajectory was recorded by both the *x* and *y* coordinates of the dot and the angle at a sampling rate of 60 Hz. To define the anchors of the scale, we modified the generalized Labeled Magnitude Scale (gLMS^65^) to make participants use the entire space of the scale. The anchors consist of no sensation (0°), weak (18°), moderate (50°), strong (108°), and very strong (162°), and strongest imaginable sensation (180°; see **Fig. 1d** and **Supplementary Fig 8**).

### Experimental procedure

The entire experiment unfolded over two-day sessions. In the first day session, we executed the pain calibration task to induce similar levels of pain experience across participants^8, 15^. During the second day session, we conducted the fMRI pain experiment. We utilized Psychtoolbox (http://www.psychtoolbox.org) and Matlab (MathWorks) to present stimulus, record ratings, and administer heat stimulation. A more detailed experimental procedure is described in each task section below.

### Day 1: Pain calibration task

Upon arriving at the laboratory, participants were informed about the procedures of the entire experiment and provided written informed consent. After completing a battery of individual difference measures—including demographic information, behavioral tendency, emotional states, and traits (not analyzed in this study)—we conducted the pain calibration task. The task had three objectives: 1) to familiarize participants with the heat stimulation and rating procedure, e.g., the use of a joystick and rating scales; 2) to match the levels of subjective pain experience across individuals; and 3) to exclude individuals with either excessively low or high sensitivity to heat stimulation.

In the pain calibration task, participants were asked to report their pain intensity ratings for twelve heat stimulations. Three predetermined heat temperatures (43.4, 45.4, and 47.4°C) were delivered in a pseudo-random order first, and based on the pain ratings for the three stimuli, we fitted the regression line of pain ratings on temperature levels. Then, the temperature levels of subsequent trials were determined based on the linear regression line ^8, 15^. For example, the first linear regression estimation was performed with the first three heat stimuli and corresponding pain ratings. Based on the linear regression result, three temperatures of heat stimulus were determined for the low, medium, and high levels of stimulus intensity, which corresponded to 30% (i.e., 54°), 50% (90°), and 70% (126°) of the semi-circle rating scale, between 40°C and 49.2°C in 0.2°C increments. Then, one of the three stimulus intensity levels was delivered in a pseudo-random order. This procedure was repeated every trial until the end of the task (12 trials). Then, the final five temperatures that corresponded to five stimulus intensity levels (30, 40, 50, 60, and 70% of the scale, which were 54°, 72°, 90°, 108°, and 126°, respectively) were determined based on the final linear regression model of each participant (see **Fig. 1b**). These calibrated five temperature levels for each participant were expected to deliver similar levels of pain intensity during the fMRI task.

If the estimated five temperatures fell outside the pre-defined temperature range, which spanned from 40 to 49.2°C, or if the *R*^2^ of the final linear regression model was lower than 0.4, we excluded the participants from further experimentation. In addition, based on the *R*^2^ for each skin site, we selected the top three skin sites, which were used in the subsequent fMRI experiment. **Supplementary Fig. 1** shows the pain calibration task results of the remaining participants. For the calibration task, participants used a joystick to report pain ratings, which helped them become familiar with the setting for the fMRI experiment. The lighting and temperature of the room in which the pain calibration task was conducted were maintained similarly to those in the MRI room. To administer heat stimulations to the same skin sites in the subsequent fMRI task, we photographed the left forearm of each participant after completing the pain calibration task.

### Day 2: fMRI experiment

On day 2, we conducted an fMRI experiment, which included one resting-state run, two simple motor task runs, six pain prediction task runs, and a structural scan (T1). First, we acquired structural brain images. During the structural scan, we also administered a pain prediction task with eight runs to help participants become familiar with the task. In addition, we provided cues consistent with the subsequent stimulus intensity to enhance the cue-induced expectation effects. During this practice, heat stimulations were delivered to a skin site that was not selected from the pain calibration task. Second, we conducted a resting-state run (1 run), lasting approximately 6 minutes and 10 seconds (810 TRs). Participants were instructed to gaze at a fixation cross on a dark gray screen. Third, we administered a simple motor task (2 runs), wherein participants were instructed to move the orange-colored dot to a specific location on a semi-circular rating scale using an MR-compatible joystick. The simple motor task runs were positioned prior to the first and fourth pain prediction task runs.

Fourth, we conducted the continuous pain prediction task (6 runs). Each run comprised 18 trials (i.e., a total of 108 trials), and each trial included a series of the following events: 1) pain-predictive cues, 2) pain prediction rating and heat stimulation, and 3) overall pain rating. More specifically, during the cue event, twenty-five dots on the semicircular rating scale were displayed prior to heat stimulation. Participants were informed that the dots represented other participants’ pain ratings for the upcoming stimulus. Prior studies showed that no additional learning procedure was needed to learn these social information-based cues, thereby simplifying our experimental procedure ^13, 32^. In reality, we generated twenty-five dots by sampling values from a normal distribution with varying means and a standard deviation (mean = 39.6° and 138.6 ° corresponding to ‘low pain cue’ and ‘high pain cue’ respectively, with a standard deviation = 9°). Some trials proceeded without cues (the ‘no cue’ condition; see **Fig. 1b**).

During the pain prediction rating and heat stimulation event, a rating scale with multiple semi-circles and an orange-colored dot at the scale’s center was displayed 1 second before the delivery of the heat stimulus. Participants were instructed to continuously report their ratings in response to the following question, “How much pain do you predict?” We informed participants that they were not required to reach the outer end of the semi-circles and that the angle, not the distance from the starting point, reflected their current pain prediction. In addition, we instructed participants to start reporting their pain prediction as soon as the trial began, ensuring they provided their ratings throughout the entire trial. We imposed a limit on the speed of the dot movement to ensure participants did not reach the outer end of the semi-circles before the heat stimulus reached its plateau.

During the overall pain rating event, participants were instructed to report their ratings in response to the following questions, “How painful was it?” (self-pain question) or “How painful would it be for others?” (other-pain question) within five seconds. Even if the participant reported pain before the end of the five seconds, the blank screen remained until the end of the five seconds. Within a run of 18 trials, the self-pain question was asked for 11 trials, and the other-pain question for 7 trials. The 11 trials with the self-pain question included all combinations of stimulus intensity levels and cue conditions, allowing us to analyze the cue effects on the overall pain ratings (**Fig. 1c**). A fixation cross was displayed during the jittered inter-trial and inter-stimulus intervals, which lasted between 3 and 7 seconds, totaling 15 seconds for each trial (see **Fig. 1a**). The trial sequence was pseudo-randomized to prevent the same experimental condition from occurring consecutively.

### fMRI data acquisition and preprocessing

Both functional and structural images were obtained using a Siemens 3.0 Tesla Magnetom Prisma at the Center for Neuroscience Imaging Research in Sungkyunkwan University utilizing a 64-channel head coil. High-resolution anatomical T1-weighted images were acquired using the MPRAGE protocol (TR = 2,400 ms; TE = 2.34 ms; flip angle = 8; field of view = 224 mm; voxel size = 0.7 mm). We acquired functional images using the T2*-weighted echo-planar image (EPI) protocol (TR = 460 ms; TE = 27.20 ms; multi-band factor = 8; field of view = 220 mm; flip angle = 44°; the number of interleaved slices = 56; voxel size = 2.7 mm^3^). For all functional scans, we discarded the initial 18 TR images to allow for image intensity stabilization. Prior to statistical analyses, we implemented the following preprocessing steps: outlier detection, realignment, distortion correction, coregistration and normalization, and spatial smoothing. 1) Outlier detection: outliers were identified in each image based on Mahalanobis distances and the root mean square of successive differences to eliminate intermittent gradient and severe motion-related artifacts. These types of artifacts are present to some degree in all fMRI data. For Mahalanobis distance-based outlier detection, we computed the distances for the matrix of concatenated slice-wise means and SDs over time and then identified images that exceeded 10 mean absolute deviations based on moving averages with a full width at half maximum (FWHM) of 20 image kernels as outliers. Using the root mean square of successive differences across volumes, images exceeding three SDs from the global mean were identified as outliers. Time points flagged as outliers by either detection method were included as nuisance covariates. 2) Realignment: functional images were realigned to the first single-band reference (SBRef) images, and six movement parameters (*x*, *y*, *z*, *pitch*, *roll*, and *yaw*) were estimated for each run. 3) Distortion correction: susceptibility-induced image distortion was corrected using TOPUP tool in the FMRIB Software Library (FSL). 4) Coregistration and normalization: we first coregistered the T1 structural images to the first SBRef functional image, segmented and normalized the structural images to the standard brain template, which was the MNI space, and then normalized the functional images to the MNI space using the parameters obtained from the T1 normalization with the interpolation to 2 mm^3^ voxels. 5) Smoothing: we smoothed the normalized functional images with a 5-mm full-width-at-half-maximum Gaussian kernel. Except for the distortion correction, the functional and structural data were preprocessed using Statistical Parametric Mapping (SPM12; Wellcome Trust Centre for Neuroimaging). The fMRI data of one participant was excluded due to an unexpected technical error. Thus, the remaining 58 participants’ data were further analyzed.

### Continuous rating data preprocessing

The sampling rate of the rating trajectory was 60 Hz, which was the refresh rate of a display projector (Propixx, VPixx Technologies; resolution of 1920 × 1080 pixels). However, due to occasional instability in the refresh rate, the number of data points was not identical across trials. Thus, we resampled the ratings to 50 Hz using linear interpolations ^21^, yielding 726 data points for the pain prediction ratings, and applied the gaussian smoothing (window size = 71) to filter out the irregularities such as noise or jitter of the rating trajectory. We then segmented the continuous pain prediction ratings into 32 time-bins and averaged the ratings for each bin. This resulted in each time bin being 460 ms, aligning with the temporal resolution of our fMRI data. Lastly, we normalized the angle, which ranged from 0° to 180°, to a scale from 0 to 1. For the overall pain ratings, we also resampled the ratings to 50 Hz using linear interpolations and averaged the rating trajectories at the last 1 sec rating as the overall pain ratings.

### Multilevel general linear model (GLM) analysis

To evaluate the effects of the pain-predictive cues and the stimulus intensity on ratings, we performed multilevel GLM analyses based on the trial-level behavioral data. In the GLM predicting the overall pain ratings, the cue conditions were coded as -1, 0, and +1, corresponding to low pain cue, no cue, and high pain cue, respectively. The five levels of stimulus intensity were coded as 1, 2, 3, 4, and 5. Additionally, an interaction term between the cue and stimulus intensity was included in the model. We used the averaged ratings over the last 1 second of the overall pain rating trajectory for the self-pain question trials. For significance testing, we conducted bootstrap tests with 10,000 iterations.

In the GLM predicting the continuous pain prediction ratings, we used ratings from each time point as an outcome variable. Considering that the continuous ratings comprised 32 time bins, a total of 32 multilevel GLM analyses were conducted. In this analysis, to help comparisons of the effect magnitude between the cue and stimulus intensity effects, we coded the cue conditions and stimulus intensity levels based on the normalized rating scale. More specifically, the low pain cue, high pain cue, and no cue conditions were coded as 0.22 (39.6°), 0,77(138.6°), and 0, respectively, and the stimulus intensity levels 1 to 5 were coded as 0.3, 0.4, 0.5, 0.6, and 0.7, respectively. This is the same as the positions of stimulus intensity levels and the mean positions of cues on the semi-circular rating scale. For significance testing, we conducted bootstrap tests with 10,000 iterations and corrected the results for multiple comparisons using Bonferroni correction.

### fMRI single-trial analysis

We estimated single-trial heat-evoked brain activity using a general linear model design matrix with separate regressors for each trial, as in the “beta series” approach^66^. In this analysis, we modeled the heat stimulation epoch of each trial with a boxcar function regressor and then convolved it with a canonical hemodynamic response function. We also included additional event regressors, including cue presentation and overall pain rating, and the nuisance covariates, including dummy coding regressors for each run (i.e., run intercept), linear drift across time within each run, indicator vectors for outlier time points, 24 head motion parameters, including six movement parameters (*x*, *y*, *z*, *pitch*, *roll*, and *yaw*), their mean-centered squares, their derivatives, and squared derivatives, and five principal components of cerebrospinal fluid and white matter signal. In addition, we applied a high-pass filter with a cutoff of 180 seconds. As single-trial estimates could be affected by acquisition artifacts occurring during that trial (for example, sudden motion, scanner pulse artifacts, etc.), we calculated trial-by-trial variance inflation factors (VIFs; a measure of design-induced uncertainty due to collinearity with nuisance regressors) using the design matrix. Any trials with VIFs that exceeded 2.5 were excluded from subsequent analyses ^10, 13, 17^. The average number of excluded trials was 3.51 (approximately 3%) with a standard deviation (SD) of 2.90.

### fMRI single-trial finite impulse response (FIR) analysis

We also conducted the single-trial FIR analysis. Instead of using a boxcar regressor for each trial, we employed a stick function, convolved with a canonical hemodynamic response function, as a regressor for each TR. We modeled a total of 45 TRs for each trial, which equated to 20.7 seconds (= 45 TRs × 0.46 seconds) from the onset of the heat stimulation. Other nuisance covariates remained consistent with the fMRI single-trial model described above.

### Whole-brain multilevel mediation analysis

To search for the brain regions mediating the effects of pain-predictive cues or stimulus intensity on the overall pain ratings, we conducted a whole-brain search for significant mediators with the multilevel mediation models. We tested the same two models described in the previous section with each voxel’s brain activity as a mediator. We conducted bootstrap tests with 10,000 iterations for significance testing and corrected the results for multiple comparisons using false discovery rate *q* < 0.05 ^67^. For more details of the multi-level mediation analysis, please see refs.^15, 17, 68^.

### Whole-brain multilevel temporal mediation analysis

We performed a whole-brain multilevel temporal mediation analysis to investigate the temporal dynamics of brain mediation for the cue and stimulus effects on the continuous pain prediction ratings. In this mediation model, Path *a* represents the effects of experimental conditions (*x*) on heat-evoked brain activity (*m*) at each time point. Path *b* is the effects of brain activity (*m*) on continuous pain prediction ratings (*y*) for a specific TR time bin above and beyond the effects of experimental conditions. Path *a*×*b* is the mediation effects, defined by the product of Paths *a* and *b.* Testing the significance of Path *a*×*b* is equivalent to a statistical test of Path *c* minus Path *c’*, where Path *c* indicates the effects of experimental conditions (*x*) on pain prediction ratings (*y*), and Path *c’* represents the effects of experimental conditions (*x*) on pain prediction ratings (*y*) after controlling for the effects of brain activity (*m*) on pain prediction ratings (*y*).

The temporal mediation analysis entails testing multilevel mediation models for all combinations of 45 TR (= 20.7 seconds) heat-evoked brain activity derived from the single-trial finite impulse response (FIR) model and 32 segmented continuous pain prediction ratings (= 14.5 seconds). We conducted the analysis for each combination separately and then combined the results, creating 45 × 32 result matrices (see **Supplementary Fig. 4a**). Brain data were resampled to a resolution of 4 mm^3^ using FLIRT in FSL to reduce computational load, resulting in 24,860 voxels for the whole brain. We also applied temporal smoothing with a Gaussian kernel using smoothdata.m function in Matlab (MathWorks) with a window size of 5. We eliminated uninterpretable time domains from further analyses, including the mediation effect of later brain activity on earlier ratings or the influence of stimulus intensity on ratings preceding thermal stimulation. These excluded time domains appear as blank in the result matrices depicted in **Supplementary Fig. 4a**.

### Independent component analysis

To identify temporal domains for brain mediation, we employed independent component analysis (ICA). In general, the ICA allows us to decompose complex patterns of data into statistically independent components. Here ICA was used to identify independent temporal domains and corresponding brain mediators, revealing the mediation effects of certain brain regions on pain prediction ratings at a specific time domain. The temporal mediation result matrices, specifically *p*-values for Path *a*×*b*, served as the input for the ICA. To make the input matrix, we first vectorized the mediation result matrices (i.e., *p*-values for Path *a*×*b*) for each voxel and converted them into -log_10_(*p*). Next, we concatenated these vectors across all voxels, thereby aggregating the data from the whole brain. We then obtained temporal and spatial component weights using the GIFToolbox with the fastICA algorithm (# of components = 5).

The analysis yielded two sets of five components, one for spatial weights (24,860 [# of voxels]× 5 [# of components]) and the other for temporal weights (880 or 602 [# of temporal mapping] × 5 [# of components]) (see **Supplementary Fig. 4a**). To identify the brain mediators and relevant temporal domains for each component, we first thresholded the temporal weights with the top 2.5 percentile. The resulting five thresholded temporal weights could be described using a river plot connecting the brain activity and pain ratings (**Supplementary Fig. 4a**). We then identified brain mediators related to these temporal domains based on the following three criteria: 1) voxels that survived the FDR correction for multiple comparisons at *q* < 0.05, 2) regions with at least five contiguous voxels, and 3) the survived voxels should cover at least 5% of the defined temporal domain. The resulting thresholded ICA component weights can be found in **Supplementary Figs. 4b-c**. One of the identified components was excluded from further analysis due to its strong association with the visual network, potentially reflecting task-related processes. For a visualization purpose, brain mediators were resampled at 2 mm^3^ and spatially smoothed with a 2-mm full-width-at-half-maximum Gaussian kernel.

### Principal gradient analysis

To further interpret the brain mediation results based on a cortical hierarchy, we first derived the functional gradient map (only the first gradient) using a resting-state fMRI from the current study participants (*N* = 56). The gradient map was constructed using BrainSpace Toolbox^69^ (see *Supplemental Methods* and **Supplementary Fig. 9**). We estimated the principal gradient using the method employed by ref. ^24^. We divided the principal gradient into 10 or 100 bins (using 10-percentile and 1-percentile bins, respectively), resulting in 10 or 100 binary gradient bin masks. Next, we calculated the overlapping proportions between the supra-thresholded brain maps and the gradient bins.

### Large-scale functional network overlap analysis

The radial plots in **Figs. 4** and **5** show the relative proportions of the number of overlapping voxels between the thresholded mediation maps and each network (or region) given the total number of voxels within each network (or region). We used the Buckner group’s parcellations to define large-scale functional brain networks, including seven networks within the cerebral cortex^70^, cerebellum^71^, and basal ganglia^72^. We also added the thalamus, hippocampus, and amygdala from the SPM anatomy toolbox and the brainstem region^73^, as shown in **Supplementary Fig. 10**.

## Author Contributions

S.G. and C.-W.W. conceived, designed, and conducted the experiment. S.G. and C.-W.W. analyzed the data and wrote the manuscript. S.G., S.-J.H., E.A.R.L., and C.-W.W. interpreted the results and edited the manuscript. C.-W.W. provided supervision.

## Acknowledgments

We thank Taenyun Kim, SeongJae Park, Minie Jung, Jinwon Park, Hong Ji Kim, and Soo Ahn Lee for their help in conducting the experiment. This work was supported by IBS-R015-D1 (Institute for Basic Science; to C.-W.W.), 2021M3E5D2A01022515 (National Research Foundation of Korea; to C.-W.W.), and HI19C1328 (Korea Health Technology R&D Project through the Korea Health Industry Development Institute (KHIDI), the Ministry of Health & Welfare, Republic of Korea; to S.G.).

## Data Availability

The data used to generate figures, including the predictive models, will be shared upon publication through a Figshare repository.

## Code Availability

The codes for generating the main figures will be shared upon publication through a Figshare repository. In-house Matlab codes and other toolboxes are available at https://github.com/canlab/CanlabCore and https://github.com/cocoanlab/cocoanCORE (for behavioral and fMRI data preprocessing, statistical analyses, and data visualization), https://github.com/canlab/MediationToolbox (multilevel mediation), https://trendscenter.org/software/gift/ (ICA), https://www.fil.ion.ucl.ac.uk/spm/software/spm12/ (SPM12), https://fsl.fmrib.ox.ac.uk/fsl/fslwiki (FSL), https://github.com/MICA-MNI/BrainSpace (BrainSpace toolbox), and https://www.mccauslandcenter.sc.edu/mricrogl/ (MRICroGL).

