## Supplementary Information for "Temporal Dynamics of Brain Mediation in Predictive Cue-induced Pain Modulation"

**This PDF file includes:**

Supplementary Methods (pp. 2-3)
Supplementary Figures 1-10 (pp. 4-15)
Supplementary References (p. 16)

### SUPPLEMENTARY METHODS

#### ROI-based temporal mediation analysis

We conducted a region-of-interest (ROI)-based approach to complement the results of the whole-brain temporal mediation analysis results. This approach allows us to examine the temporal dynamics of brain mediation for specific regions, minimizing the need for some exploratory approaches such as ICA. Based on prior literature[1-3], we selected 17 *a priori* ROIs known for their roles in pain processing. These include areas involved in nociceptive pain processing[1] (e.g., left and right thalamus, right second somatosensory area, and left and right dorsal-posterior insula cortex), regions associated with self-regulatory strategies in response to pain[2] (e.g., ventromedial prefrontal cortex and nucleus accumbens), and areas linked to social information processing related to pain[3] (e.g., pregenual anterior cingulate cortex, orbitofrontal cortex, left and right ventrolateral prefrontal cortex, left supramarginal gyrus, medial frontal gyrus, intraparietal sulcus, and right dorsolateral prefrontal cortex) (see **Supplementary Fig. 7a**).

For the ROI-based temporal mediation analysis, we first obtained the average activity for each ROI across the 45 TRs from the single-trial finite impulse response (FIR) model. We then assessed the mediation effects of ROI activity at specific time points on continuous pain prediction ratings (spanning 32 TR bins). In our mediation model, cue conditions (coded as -1,0, and 1) and stimulus intensity levels (coded as 1 to 5) were the predictors ($x$), ROI activity served as the mediator ($m$), and pain prediction ratings were the outcome variable ($y$). When cue condition served as a predictor, stimulus intensity was included as a covariate, and in the model where the stimulus intensity served as a predictor, the cue levels were included as a covariate.

To test the significance of the mediation effects, we employed bootstrap tests with 10,000 iterations. This process resulted in 45×32 result matrix for each ROI, similar to the approach taken in the whole-brain analysis. To identify the significant temporal domains within each ROI, we used the "spm_bwlabel.m" function in SPM12 (Wellcome Trust Centre for Neuroimaging). This function detects connected multiple clusters within a 2D temporal domain result matrix after applying a threshold of $q < 0.05$, false discovery rate corrected (FDR)[4]. The results of this analysis are shown in **Supplementary Fig. 7b-c**.

#### Building a volumetric gradient map using a resting-state fMRI data

Our study opted for a volume-based fMRI analysis approach, contrasting with the surface-based methods used in the original functional gradient study[5]. This discrepancy made it difficult to apply the publicly available functional gradient map to our data directly. To circumvent this problem, we created our own volumetric gradient map using resting state-fMRI data from the participants of the current study ($N = 56$) using the BrainSpace toolbox[6]. From our original 59 participants, we had to exclude three participants from this analysis due to technical issues, such as preprocessing errors, resulting in $N = 56$. To create the volumetric gradient map, we first resampled the resting-state fMRI data to a voxel size of 3-mm$^3$ to reduce the computational load to a manageable level. We then computed the functional connectivity based on the resting-state fMRI activity with the gray matter mask. Finally, the functional connectivity matrices from 56 participants were averaged, resulting in a single averaged connectivity matrix with a size of 59,026 × 59,026 (voxel-by-voxel). This averaged connectivity matrix was then submitted to the BrainSpace toolbox[6] with the same parameters as the original study[5] (i.e., dimension reduction technique: diffusion embedding, kernel: normalized angle, sparsity: 0.9; see **Supplementary**

**Fig. 9**).

### SUPPLEMENTARY FIGURES

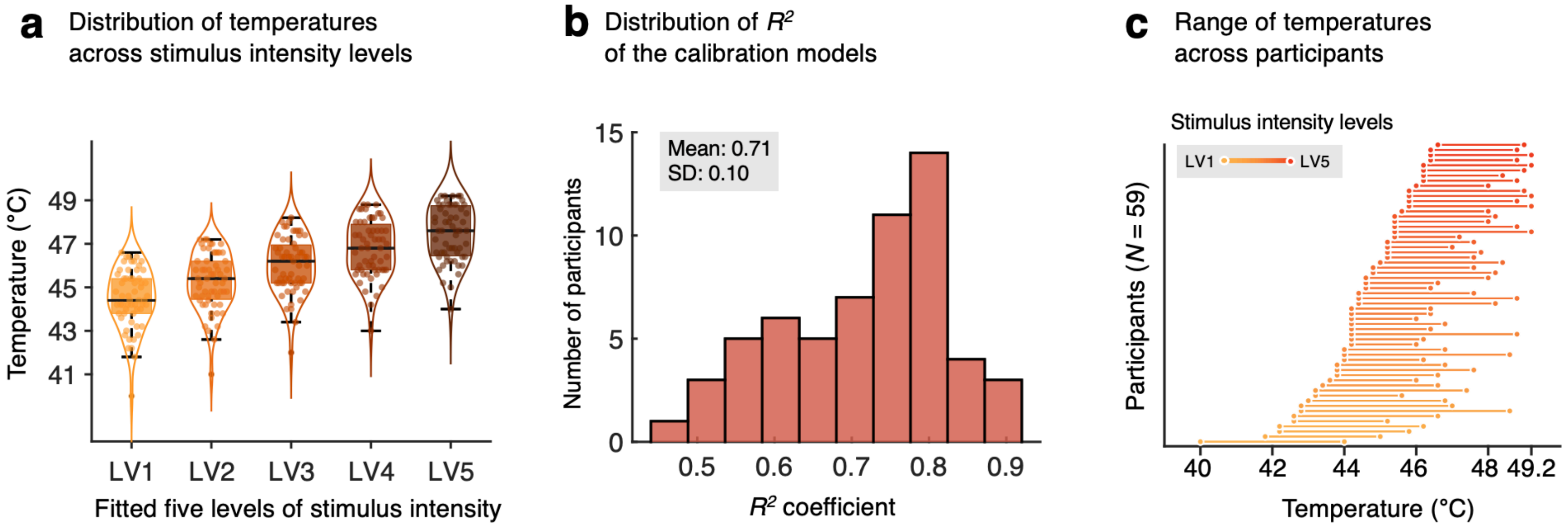


**Supplementary Fig. 1. Results of pain calibration task. a,** The pain calibration task was conducted to determine individualized stimulus intensity levels to match the levels of subjective pain experience across individuals. Each dot represents the temperature calibrated for an individual, corresponding to each stimulus intensity level. The mean temperatures (standard deviation) for each stimulus level are as follows: 44.49°C (1.34) for LV1, 45.25°C (1.30) for LV2, 46.02°C (1.27) for LV3, 46.78°C (1.29) for LV4, and 47.47 °C (1.26) for LV5. One-way analysis of variance results revealed a significant main effect of the fitted stimulus intensity, $F$ (4, 290) = 49.02, $p$ = 1.786e-31. **b,** The distribution of the $R^2$ coefficients of the final linear regression models from the pain calibration task. **c,** Each line represents the range of calibrated stimulus intensities for each participant, and dots at the end of lines indicate LV1 and LV5 stimulus intensity. The order of participants is sorted by the temperature of LV1. Data from 59 participants who passed the pain calibration task were displayed in this figure.

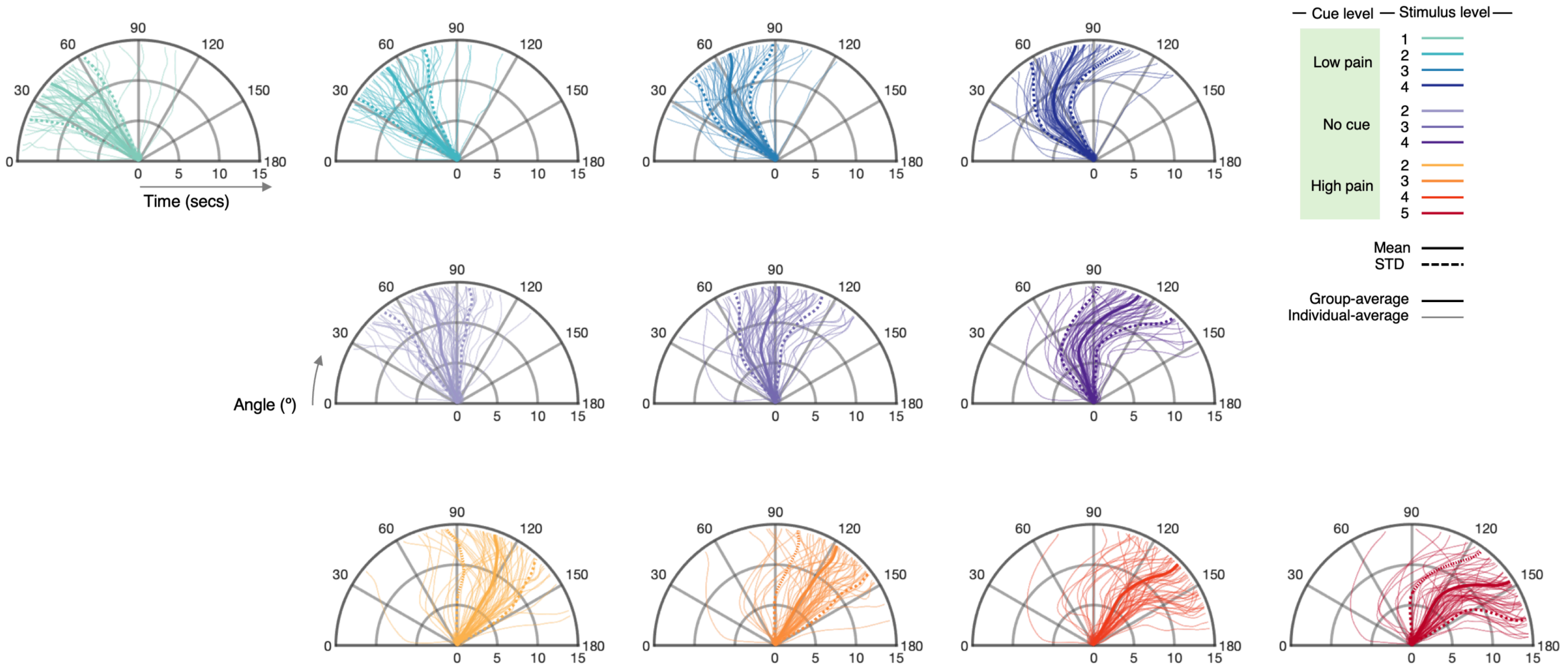


**Supplementary Fig. 2. Average trajectory of continuous pain prediction ratings for all experimental conditions.** Thick and solid lines represent the group average of rating trajectory, while dotted lines represent the within-subject standard error of the mean (s.e.m.). Thin lines illustrate individual averages of rating trajectories. Different colors correspond to experimental conditions.

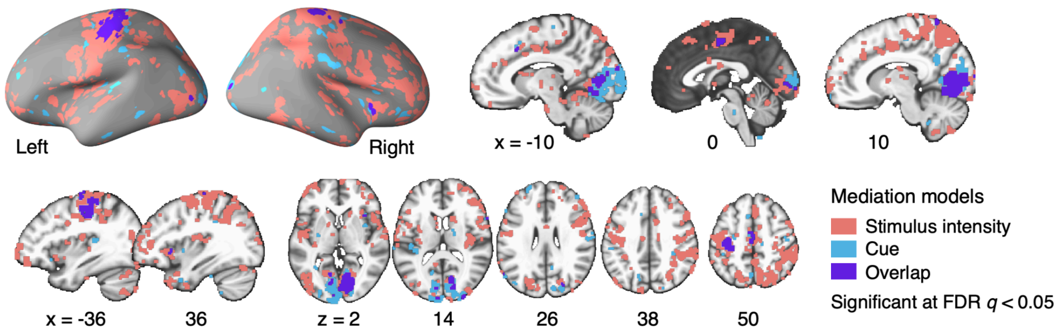


**Supplementary Fig. 3. Conjunction of the cue and the stimulus mediation maps.** The conjunction map shows the spatial overlap and unique areas involved in the cue and stimulus mediation. Regions colored in cyan and pink denote brain areas associated with the mediation of cue and stimulus effects, respectively. These are thresholded at voxel-wise FDR $q < 0.05$ with cluster extent $k > 5$. The areas where these two mediation maps overlap are colored in purple.

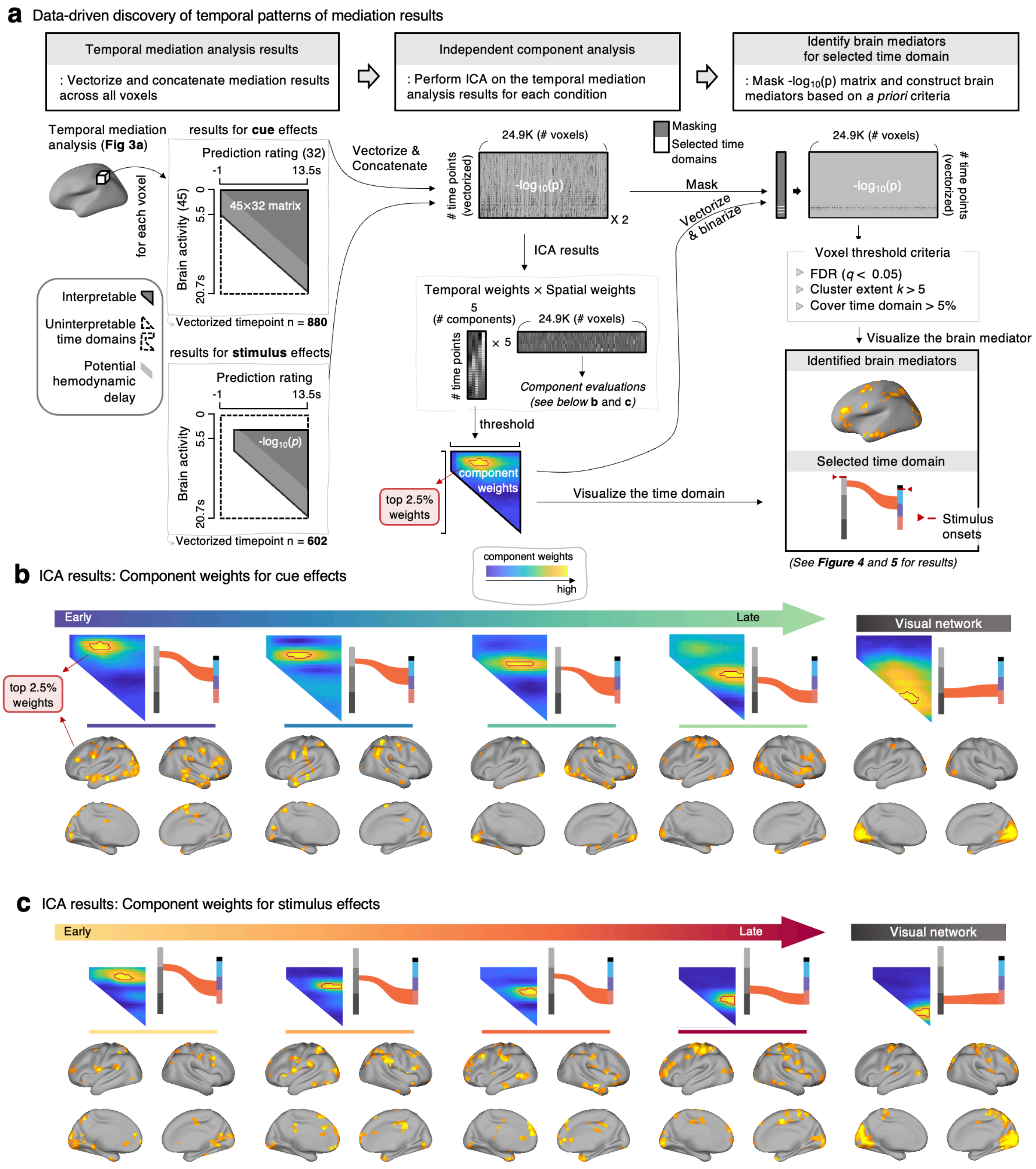


**Supplementary Fig. 4. Independent Component Analysis scheme for the temporal mediation analysis results.**

**a,** A schematic diagram that illustrates the step-by-step procedure to identify the temporal domains and brain mediators through independent component analysis (ICA). The temporal mediation result matrices ($p$-values for Path $a$×$b$) served as the input for the ICA. We eliminated uninterpretable time domains from analyses, including the mediation effect of later brain activity on earlier ratings or the influence of stimulus intensity on ratings preceding thermal stimulation. These excluded time domains appear as blank in the result matrices. To make the input matrix, we first vectorized the mediation result matrices (i.e., p-values for Path $a \times b$) for each voxel and converted them into -log10($p$). Next, we concatenated these vectors across all voxels, thereby aggregating the data from the whole brain. This resulted in matrices with dimensions of 880 or 602 by 24,860. We then obtained temporal and spatial component weights using the GIFToolbox with the fastICA algorithm (# of components = 5).

The analysis yielded two sets of five components, one for spatial weights (24,860 [# of voxels]× 5 [# of components]) and the other for temporal weights (880 or 602 [# of temporal mapping] × 5 [# of components]). To identify the brain mediators and relevant temporal domains for each component, we first thresholded the temporal weights with the top 2.5 percentile. The resulting five thresholded temporal weights could be described using a river plot connecting the brain activity and pain ratings. We then identified brain mediators related to these temporal domains based on the following three criteria: 1) voxels that survived the FDR correction for multiple comparisons at $q < 0.05$, 2) regions with at least five contiguous voxels, and 3) the survived voxels should cover at least 5% of the defined temporal domain.

**b-c,** The matrices and brain maps represent the spatial weights and temporal components from the ICA analysis. Warm vs. cool colors represent high vs. low component weights. The areas enclosed by the red outlines represent the top 2.5% of the weights. One of the identified components was excluded from further analysis due to its strong association with the visual network (right panel), potentially reflecting task-related processes.

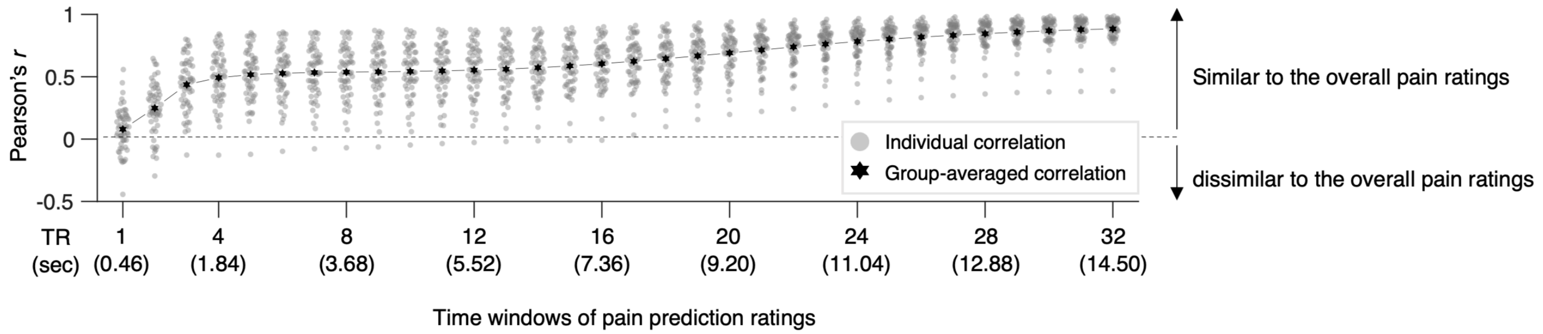


**Supplementary Fig. 5. Correlations between the continuous pain prediction ratings and the overall pain ratings.** To investigate the relationship between continuous pain prediction ratings at each time window (comprising 32 TR-level bins) and overall pain ratings, we averaged the continuous pain ratings within each TR window. We then calculated Pearson's correlation between these time-averaged continuous ratings and the overall pain ratings. The resulting plot demonstrates an overall increase in correlations throughout the trial. Analysis of variance (ANOVA) results revealed a significant time effect, $F(29,1740) = 62.35$, $p = 1.60276e\text{-}244$. This finding implies that the later stages of the continuous pain prediction ratings provide more information into the overall pain ratings compared to the earlier and middle stages.

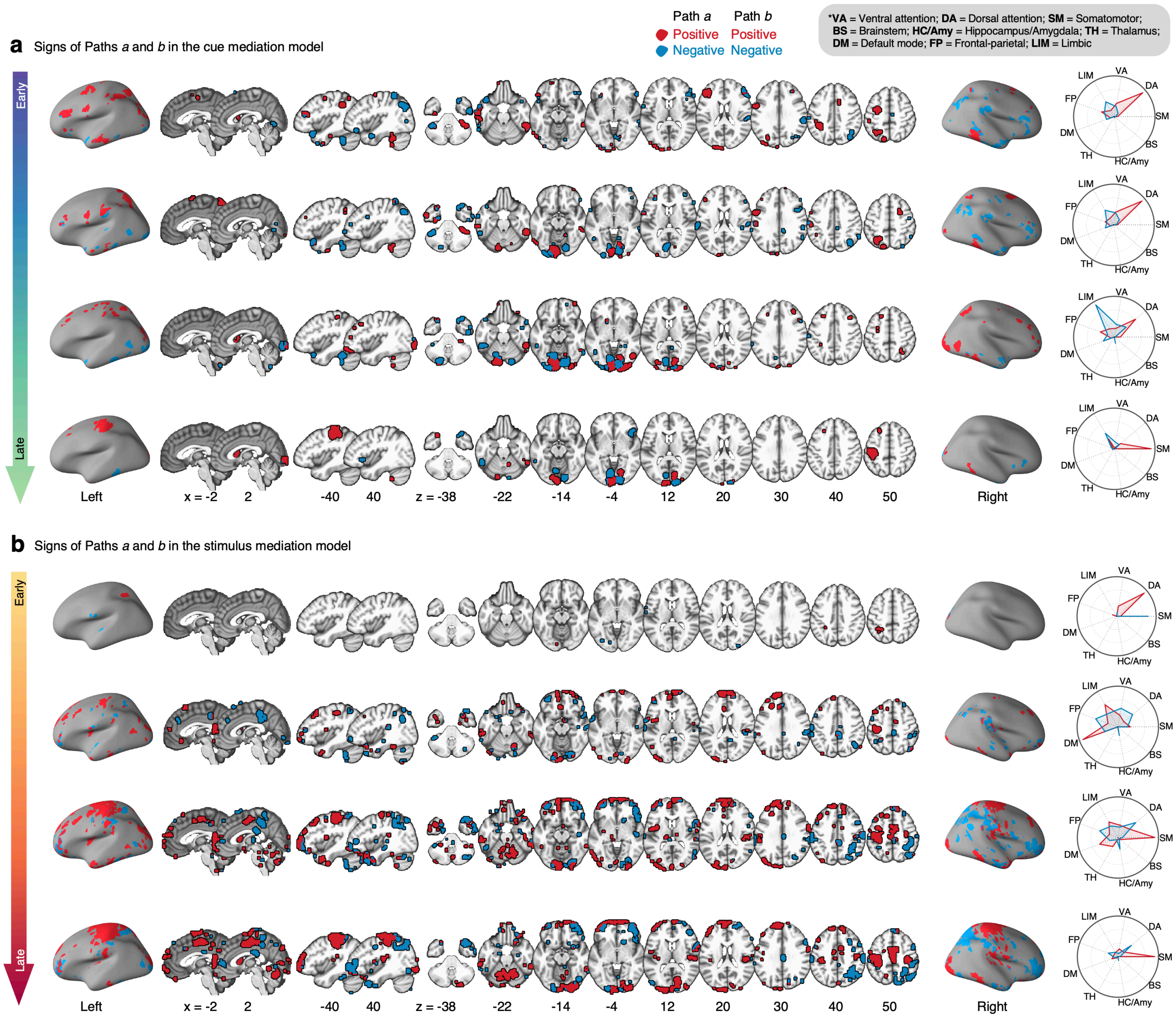


**Supplementary Fig. 6. Signs of Paths *a* and *b* of brain mediators for cue and stimulus intensity effects.** Each row of brain maps corresponds to temporal domains of the **(a)** cue and **(b)** stimulus mediation. Brain regions where Paths *a* and *b* are both positive are depicted in red, and regions where both Paths *a* and *b* are negative are depicted in blue. Radial plots show the relative proportions of the number of overlapping voxels between the thresholded mediation maps with positive (red) and negative (blue) Paths *a* and *b* and each of the large-scale networks (or regions) given the total number of voxels within each network (or region). VA, ventral attention network; DA, dorsal attention network; SM, somatomotor network; BS, brainstem; HC/Amy, hippocampus and amygdala; TH, Thalamus; DM, default mode network; FP, frontoparietal network; LIM, limbic network.

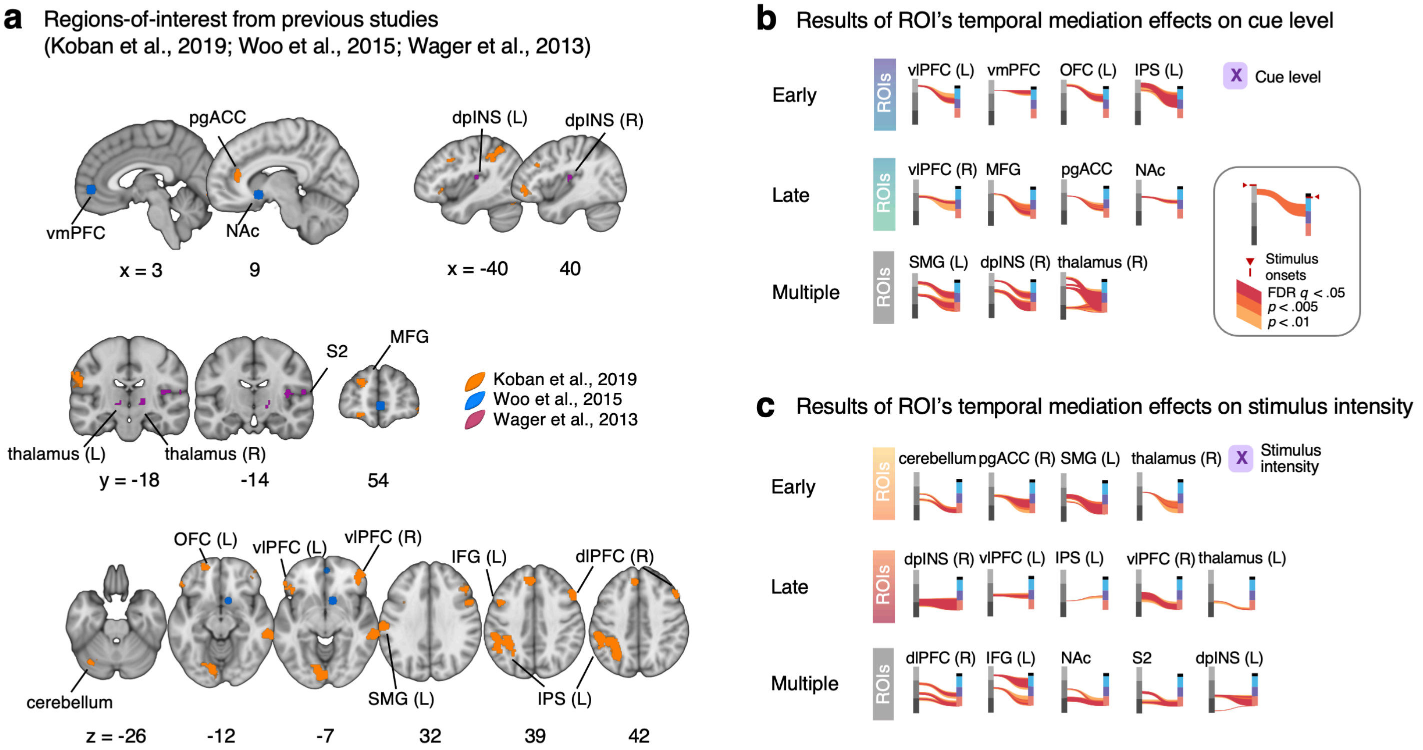


**Supplementary Fig. 7. Region-of-interest (ROI)-based temporal mediation analysis results.** We conducted a region-of-interest (ROI)-based approach to complement the results of the whole-brain temporal mediation analysis results. For a detailed description of the ROI-based temporal mediation analysis, please refer to the **Supplementary Methods**. **a,** Based on prior literature[1-3], we selected 17 *a priori* ROIs known for their roles in pain processing. These include areas involved in nociceptive pain processing[1] (e.g., left and right thalamus, right second somatosensory area, and left and right dorsal-posterior insula cortex), regions associated with self-regulatory strategies in response to pain[2] (e.g., ventromedial prefrontal cortex and nucleus accumbens), and areas linked to social information processing related to pain[3] (e.g., pregenual anterior cingulate cortex, orbitofrontal cortex, left and right ventrolateral prefrontal cortex, left supramarginal gyrus, medial frontal gyrus, intraparietal sulcus, and right dorsolateral prefrontal cortex). **b-c,** River plots illustrate the ROI-based temporal mediation analysis results. The results were categorized by brain mediation timing—'early' and 'late.' In addition, the regions with significant in more than two domains were categorized as 'multiple.' The plots show the temporal domains significant at false-discovery rate (FDR) $q < 0.05$, alongside adjacent time domains pruned using two more liberal thresholds ($p < 0.005$ and $p < 0.01$) to contextualize the results. Abbreviations: FDR, false-discovery rate; vmPFC, ventromedial prefrontal cortex; pgACC, pregenual anterior cingulate cortex; NAc, nucleus accumbens; dpINS, dorsal-posterior insular; S2, second somatosensory area; OFC, orbitofrontal cortex; vlPFC, ventrolateral prefrontal cortex; SMG, supramarginal gyrus; MFG, medial frontal gyrus; IPS, intraparietal sulcus; dlPFC, dorsolateral prefrontal cortex.

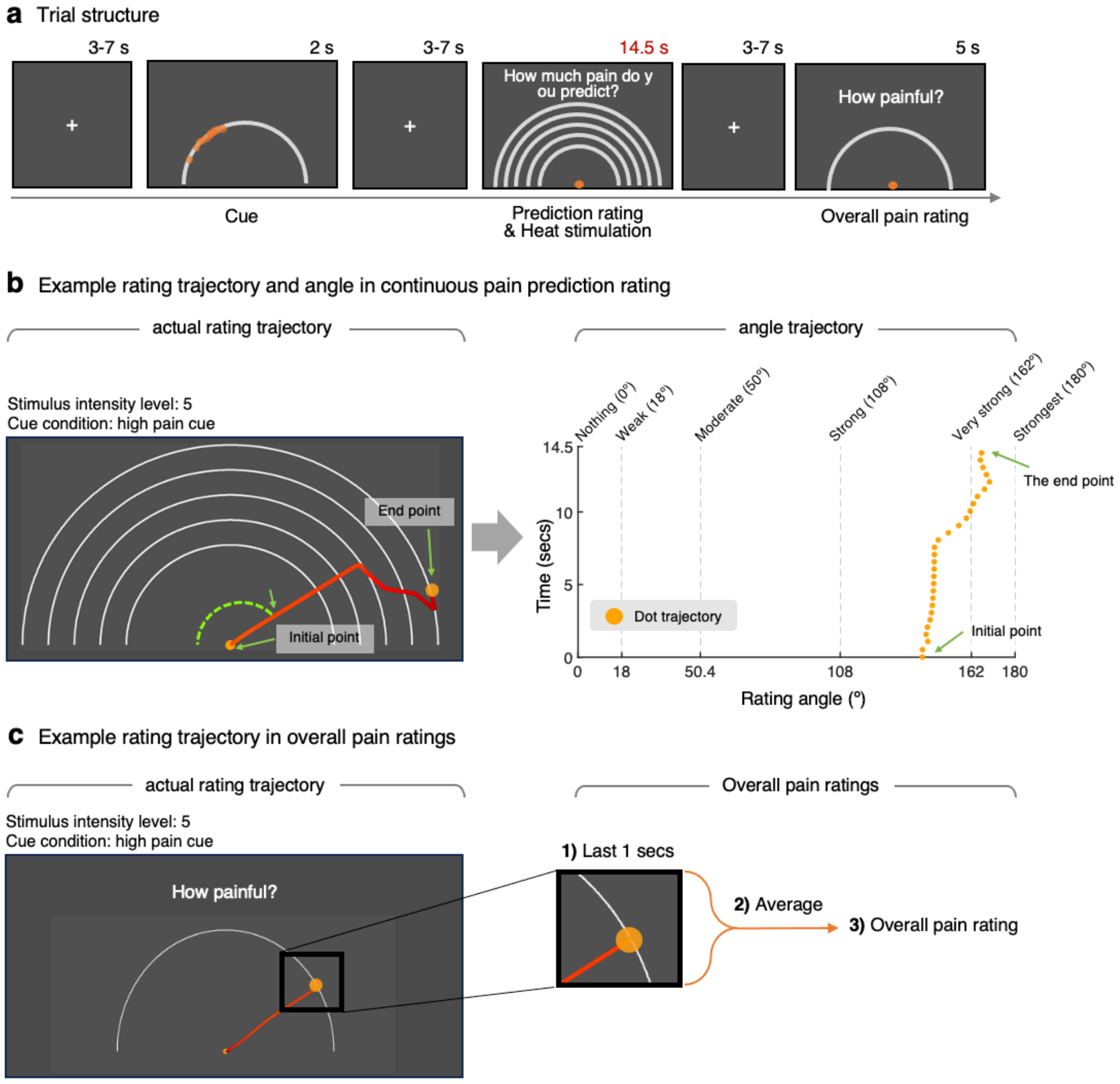


**Supplementary Fig. 8. Trial structure and rating trajectory examples. a,** Schematic overview of trial structure. **b,** Continuous pain prediction rating: (Left) The figure shows an example rating trajectory over the 14.5 seconds during the pain prediction rating period. Starting from the center (i.e., initial point), participants were asked to move the orange dot anywhere within the screen to report their continuous ratings. The participants were informed that the angle at each time point would be used as their ratings. The rating trajectory was recorded with dot's $x$ and $y$ coordinates, but it was converted to the angle from the left segment of the semicircle base. (Right) The (x, y) coordinates were converted to the angles, which served as online ratings. **c,** Overall pain rating: (Left) The figure shows an example rating trajectory for the overall pain rating. (Right) The dot's (x, y) coordinates were converted to the angles. Then, we calculated the average of the last 1 second's angles, which served as the overall pain rating.

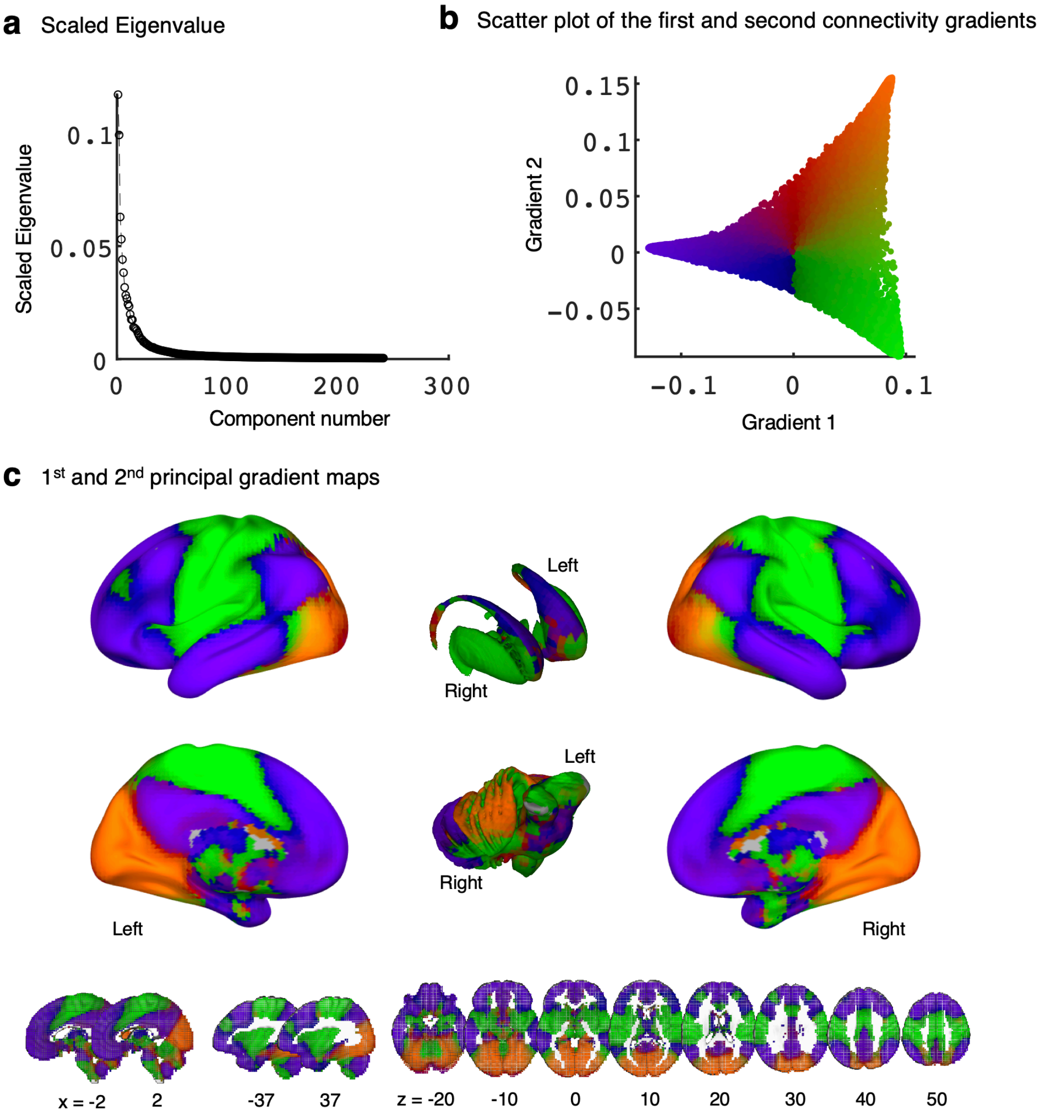


**Supplementary Fig. 9. Building a volumetric gradient map.** Principal gradient map diagnostics and resulting map. **a,** Scaled Eigenvalues across components. **b,** A scatter plot of the 1st and 2nd connectivity-based principal gradients. Each dot in the scatter plot represents a connectivity, and different color represents the relationship between the 1st and 2nd gradients. The gradient pattern was largely consistent with the original study[5]. **c,** A brain map representing the 1st and 2nd gradients using the same color scheme as the scatter plot in **b**.

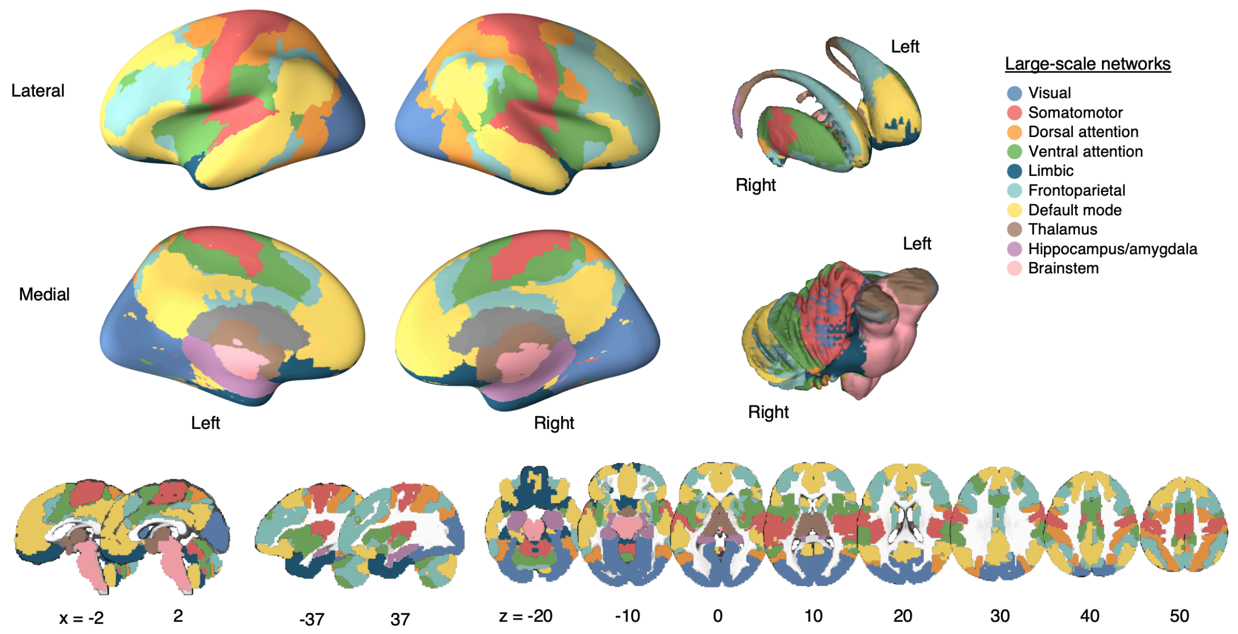


**Supplementary Fig. 10. A large-scale functional brain networks and subcortical regions.** Different colors represent different large-scale networks and subcortical regions. A large-scale functional network includes seven cortical[7], basal ganglia[8], and cerebellum networks[9]. In addition to these large-scale networks, we added thalamus, hippocampus/amygdala, and brainstem[10].
